# Divergent receptor proteins confer responses to different karrikins in two ephemeral weeds

**DOI:** 10.1101/376939

**Authors:** Yueming Kelly Sun, Jiaren Yao, Adrian Scaffidi, Kim T. Melville, Sabrina F Davies, Charles S Bond, Steven M Smith, Gavin R Flematti, Mark T Waters

## Abstract

Wildfires can encourage the establishment of invasive plants by releasing potent germination stimulants, such as karrikins. Seed germination of *Brassica tournefortii*, a noxious weed of Mediterranean climates, is strongly stimulated by KAR_1_, which is the archetypal karrikin produced from burning vegetation. In contrast, the closely-related yet non-fire-associated ephemeral *Arabidopsis thaliana* is unusual because it responds preferentially to KAR_2_. The α/β-hydrolase KARRIKIN INSENSITIVE2 (KAI2) is the putative karrikin receptor identified in *Arabidopsis*. Here we show that *B. tournefortii* differentially expresses three *KAI2* homologues, and the most highly-expressed homologue is sufficient to confer enhanced responses to KAR_1_ relative to KAR_2_ when expressed in *Arabidopsis*. We further identify two variant amino acid residues near the KAI2 active site that explain the ligand selectivity, and show that this combination has arisen independently multiple times within dicots. Our results suggest that duplication and diversification of KAI2 proteins could confer upon weedy ephemerals and potentially other angiosperms differential responses to chemical cues produced by environmental disturbance, including fire.

## INTRODUCTION

Environmental disturbance promotes the establishment of invasive species, posing a potent threat to global biodiversity. Changing wildfire regimes, such as increasing frequency of fires, is one of the most relevant disturbance factors contributing to elevated invasion threat^1^. Wildfires create germination opportunities in part by releasing seed germination stimulants, such as karrikins, from burning vegetation^2, 3^. Karrikins comprise a family of butenolides with six known members. In samples of smoke water generated by burning grass straw, KAR_1_ is the most abundant karrikin, while KAR_2_ is six times less abundant^4^, although these proportions may differ depending on source material, preparation method, age and storage conditions^5^. These two analogues differ only by the presence of a methyl group on the butenolide ring in KAR_1_, which is absent in KAR_2_ (Supplementary Fig. 1a). Invasive plant species that are responsive to karrikins could utilise natural and human-induced fires to facilitate their establishment^6, 7^.

Karrikins can overcome seed dormancy and promote seed germination in a number of smoke-responsive species, as well as others that are not associated with fire regimes^8–10^. Furthermore, karrikins also influence light-dependent seedling development. Karrikins enhance the sensitivity of *Arabidopsis thaliana* seedlings to light by inhibiting hypocotyl elongation, stimulating cotyledon expansion, and promoting chlorophyll accumulation in a dose-dependent manner^11^. Karrikins have been reported to enhance seedling survival and biomass in species such as tomato, rice and maize^12, 13^. Such growth-promoting effects of karrikins, especially at critical early stages of the life cycle, have the potential to further encourage the establishment of invasive species after fire events.

*Brassica tournefortii* (Brassicaceae; Sahara mustard) is native to northern Africa and the Middle East, but is an invasive weed that blights many ecosystems with a Mediterranean climate and chaparral-type vegetation that are prone to wildfires in North America, Australia and South Africa. *B. tournefortii* seeds can persist in the soil for many seasons, undergoing wet-dry cycling that can influence dormancy and contribute to boom-bust cycles that outcompete native species^9, 14^. *B. tournefortii* plants may radically alter fire frequency and intensity by influencing fuel profiles^15, 16^, further exacerbating the impact of fire on susceptible native ecosystems. In addition, seeds of *B. tournefortii* are particularly responsive to smoke-derived karrikins, and show a positive germination response to KAR_1_ in the nanomolar range^10^. Accordingly, *B. tournefortii* is particularly well positioned to invade areas disturbed by fire events^17, 18^.

The putative karrikin receptor KARRIKIN INSENSITIVE 2 (KAI2) was identified in Arabidopsis, a weedy ephemeral that originated in Eurasia but is now widely distributed throughout the northern hemisphere^19–21^. Arabidopsis is not known to colonise fire-prone habitats, but nevertheless seeds germinate in response to karrikins in the micromolar range^22^. Unlike most smoke-responsive species that respond more readily to KAR_1_^8, 23^, Arabidopsis responds preferentially to KAR_2_^22^. KAI2 is an evolutionarily ancient α/β-hydrolase and a paralogue of DWARF14 (D14), the receptor for strigolactones^24, 25^. Karrikins and strigolactones are chemically similar by virtue of a butenolide moiety that is necessary for bioactivity^26, 27^. KAI2 and D14 have dual functions as both enzyme and receptor, but the functional significance of the enzymatic activity remains contested^28–32^. Furthermore, the basis for ligand specificity by these two highly congruent proteins remains essentially unknown.

Orthologues of *KAI2* are ubiquitous in land plants, and are normally present as a single gene copy within an ancient and highly conserved “eu-KAI2” clade^33^. There is growing evidence that, beyond its ability to mediate karrikin responses, KAI2 has a core ancestral role in perceiving an endogenous KAI2 ligand (“KL”) that regulates seed germination, seedling development, leaf shape and cuticle development^34–36^. Since its divergence from the Arabidopsis lineage, the tribe Brassiceae, which includes the genus *Brassica*, underwent a whole genome triplication event 24–29 million years ago^37–39^. This process might have allowed additional KAI2 copies to gain altered ligand specificity, potentially enhancing perception of environmental signals such as karrikins from smoke. Here, we report that two out of three KAI2 homologues expressed in *B. tournefortii* show distinct preferences for different karrikins. We take advantage of the relatively recent genome triplication event in the Brassiceae to identify two amino acids that are sufficient to explain these karrikin preferences and confirm this by mutagenesis. Beyond demonstrating the potential ecological significance of diversity among KAI2 homologues, our findings also reveal active site regions critical for ligand selectivity among KAI2 receptor-enzymes that are found in all land plants.

## RESULTS

### *B. tournefortii* is most sensitive to KAR_1_

To characterise the karrikin response of *B. tournefortii*, we performed multiple physiological and molecular assays comparing KAR_1_ activity with that of KAR_2_. First, germination of *B. tournefortii* seeds was consistently more responsive to KAR_1_ than KAR_2_ at 10 nM, 100 nM and 1 µM (Fig. 1a and Supplementary Fig. 1b-d). Second, homologues of two karrikin-responsive transcripts, *DWARF14-LIKE2* (*BtDLK2*) and *SALT TOLERANCE HOMOLOG7* (*BtSTH7*), identified on the basis of close sequence homology to their Arabidopsis counterparts^11, 20^, were significantly more highly expressed when treated with 1 µM KAR_1_ than with 1 µM KAR_2_ (Fig. 1b). These observed differences in seed response are not due to differential karrikin uptake, since both KAR_1_ and KAR_2_ were taken up from solution at similar rates by *B. tournefortii* seeds during imbibition, as was also true for Arabidopsis seeds (Supplementary Fig. 1e-f). Besides promoting germination of primary dormant seeds, karrikins also inhibited hypocotyl elongation in *B. tournefortii* seedlings, as is the case in Arabidopsis; again, KAR_1_ showed a stronger effect than KAR_2_ (Fig. 1c-d). Levels of *BtDLK2* transcripts in seedlings were also more responsive to KAR_1_ than KAR_2_ at a given concentration (Fig. 1e). Therefore, we conclude that *B. tournefortii* is more sensitive to KAR_1_ than to KAR_2_, a ligand preference that is a feature of seed germination in many karrikin-responsive species from ecosystems prone to fires^8, 23^.

**Figure 1.**
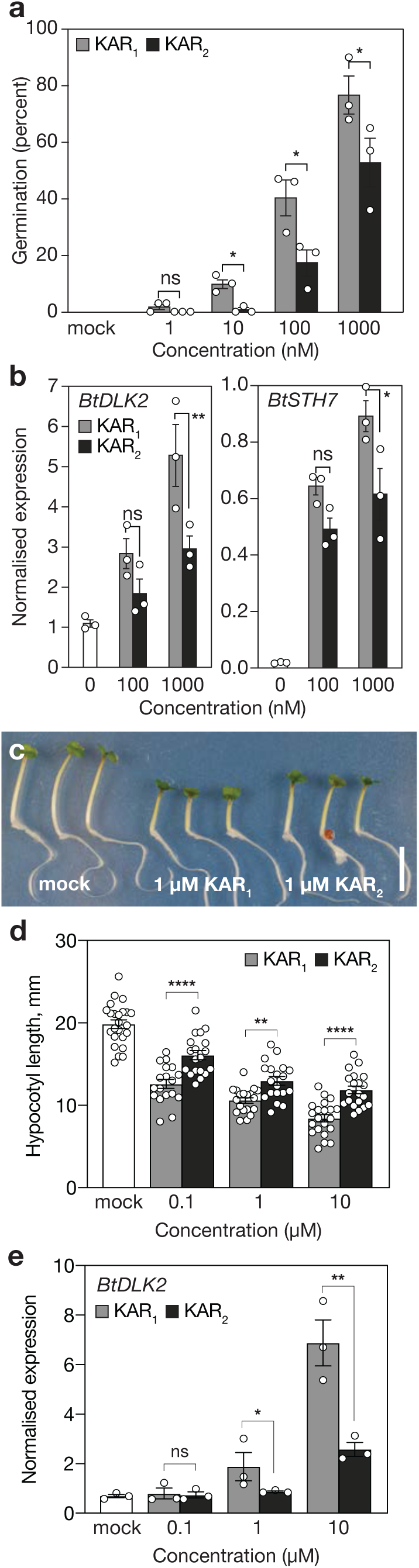
*Brassica tournefortii* is highly sensitive to KAR_1_, the major karrikin analogue isolated from plant-derived smoke. **a**, Germination responses of *B. tournefortii* seed to KAR_1_ and KAR_2_. Data are cumulative germination after 11 days (mean ± SE; n = 3 biological replicates per treatment, ≥35 seeds per replicate). **b**, Levels of karrikin-responsive transcripts *BtDLK2* and *BtSTH7* in *B. tournefortii* seed. Seed were imbibed for 24 hours in the dark supplemented with KAR_1_ and KAR_2_. Transcripts were normalised to *BtCACS* reference transcripts. Data are means ± SE, n = 3 biological replicates, ≥50 seeds per replicate. **c**, **d**, Hypocotyl elongation responses of *B. tournefortii* seedlings grown for four days under continuous red light on water-saturated glass filter paper containing 0.1% acetone (mock), KAR_1_ or KAR_2_. Data are means ± 95% CI of n = 18 to 24 seedlings. Scale bar: 10 mm. **e**, Levels of *BtDLK2* transcripts in *B. tournefortii* seedlings grown under the same conditions as for **d**. Data are means ± SE, n = 3 biological replicates, ≥20 seedlings per replicate. In all panels, asterisks denote significance levels (ANOVA) between indicated conditions: * P<0.05; ** P<0.01; *** P<0.001; **** P<0.0001. Source data are provided as a Source Data file.

### Two out of three *B. tournefortii* KAI2 homologues are functional in Arabidopsis

To establish whether there are multiple *KAI2* homologues present in *B. tournefortii*, we examined transcriptomes from seeds and seedlings. Three putative *KAI2* homologues were identified (*BtKAI2a*, *BtKAI2b*, and *BtKAI2c*; Fig. 2a; Supplementary Fig. 2 and 3). *BtKAI2a* grouped with *AtKAI2* and those of other Brassicaceae within a single clade, whereas *BtKAI2b* and *BtKAI2c* grouped within a clade unique to *Brassica*. This taxonomic distribution implies that *BtKAI2a* is most similar to *AtKAI2*, and that *BtKAI2b* and *BtKAI2c* are paralogues that arose via genome triplication during the evolution of *Brassica*. All three *BtKAI2* transcripts were expressed in *B. tournefortii* seeds, but only *BtKAI2a* and *BtKAI2b* could be detected in seedlings (Fig. 2b). In seeds and seedlings, *BtKAI2b* transcripts were the most abundant of the three, but there were no consistent effects on any of the transcripts upon treatment with 1 µM KAR_1_. We also identified two *BtD14* homologues, at least one of which is functionally orthologous to *AtD14* (Supplementary Figs. 2 and 4).

**Figure 2.**
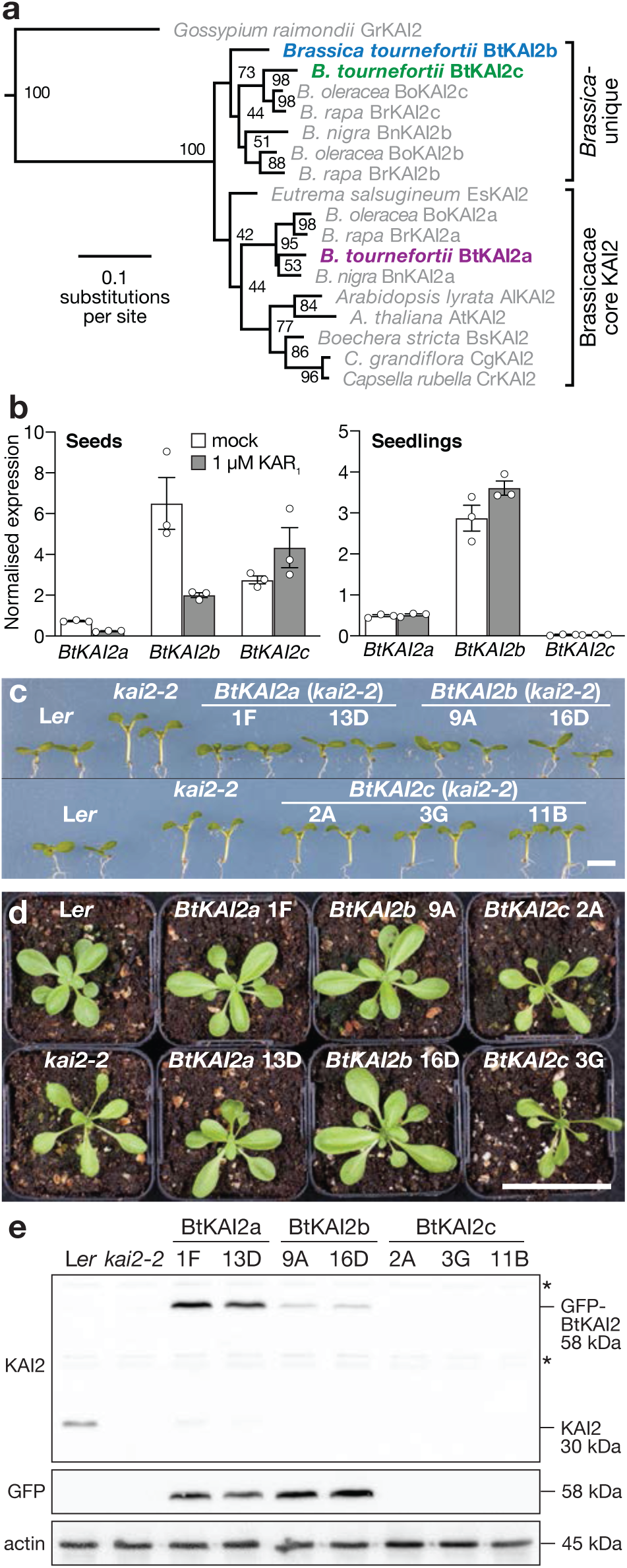
Two differentially expressed *B. tournefortii KAI2* homologues are functional in *Arabidopsis*. **a**, Maximum likelihood phylogeny of KAI2 homologues in the Brassicaceae, based on nucleotide data. Node values represent bootstrap support from 100 replicates. A eu-KAI2 sequence from *Gossypium raimondii* (Malvaceae) serves as an outgroup. Tree shown is a subset of a larger phylogeny in Supplementary Fig. 2. **b**, Transcript levels of the three *BtKAI2* homologues in *B. tournefortii* seeds imbibed for 24 h (left) and four-day-old seedlings (right) treated with 0.1% acetone or with 1 µM KAR_1_ for 24 h. **c**, **d**, Seedling and rosette phenotypes of two independent transgenic lines of Arabidopsis homozygous for *KAI2pro:GFP-BtKAI2* transgenes. Scale bars: 5 mm (**c**); 50 mm (**d**). **e**, Immunoblots of soluble proteins challenged with antibodies against KAI2 (upper panel), GFP (middle panel) or actin as a loading control (lower panel). Anti-KAI2 detects both the native AtKAI2 protein (30 kDa) and the GFP-BtKAI2 fusion proteins (58 kDa) but detects BtKAI2b relatively poorly. Non-specific bands are marked with asterisks. Protein was isolated from pools of approximately fifty 7-day-old seedlings. Source data are provided as a Source Data file.

**Figure 3.**
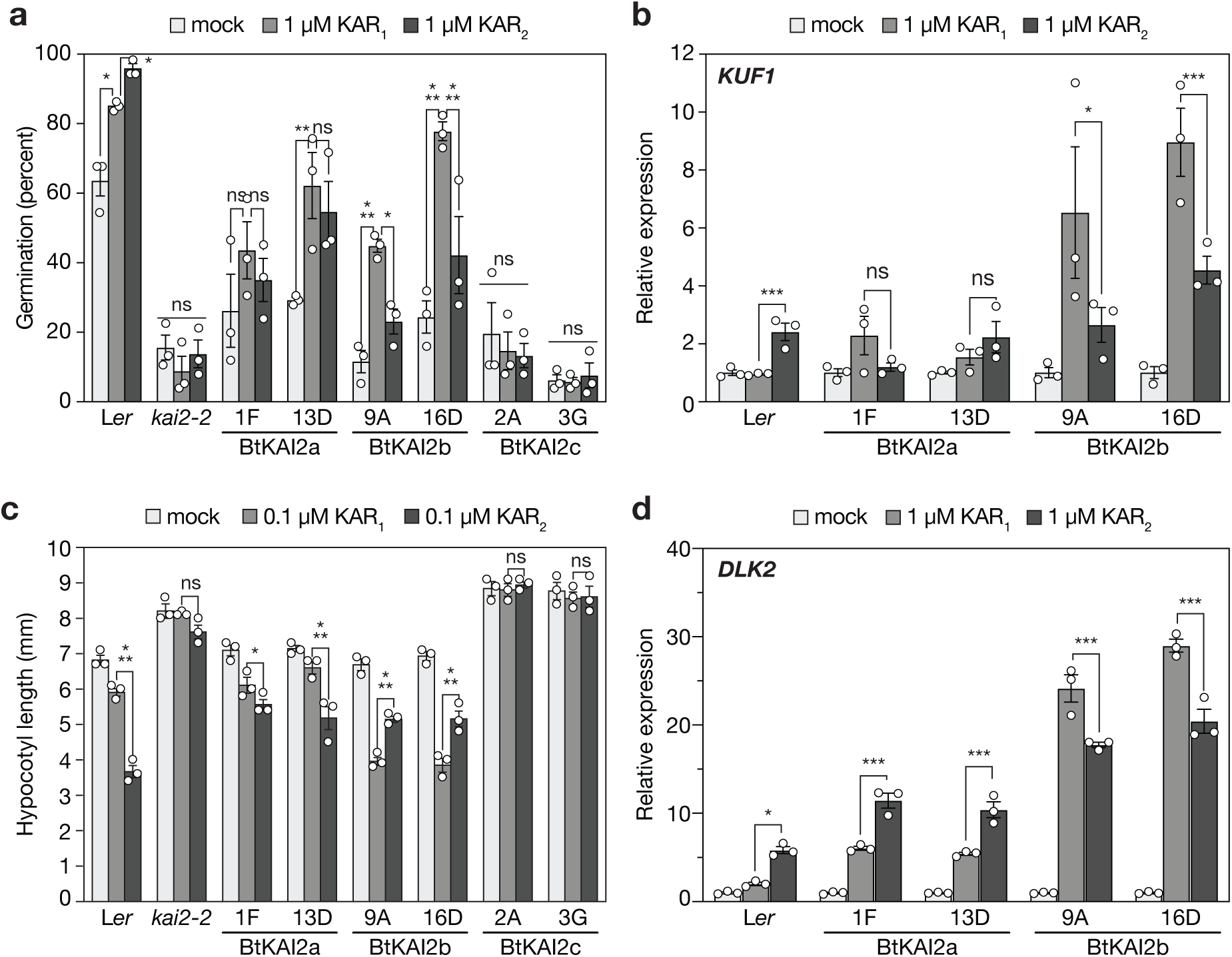
Functional divergence between BtKAI2 homologues. **a**, Germination responses of primary dormant *Arabidopsis* seed homozygous for *KAI2pro:GFP-BtKAI2* transgenes in the *kai2-2* background. Germination values were determined 120 h after sowing. Extended germination curves are shown in Supplementary Fig. 7. Data are means ± SE, *n* = 3 independent seed batches, 75 seed per batch. **b**, Levels of *KUF1* transcripts in *KAI2pro:GFP-BtKAI2* seeds treated with 1 µM KAR_1_ or KAR_2_. Expression was normalised to *CACS* reference transcripts and scaled to the value for mock-treated seed within each genotype. Data are means ± SE of n = 3 biological replicates. **c**, Hypocotyl elongation responses of *KAI2pro:GFP-BtKAI2* seedlings treated with KAR_1_ or KAR_2_. Data are means ± SE of *n* = 3 biological replicates, 12–18 seedlings per replicate. **d**, Levels of *DLK2* transcripts in 8-day-old *KAI2pro:GFP-BtKAI2* seedlings treated with KAR_1_ or KAR_2_ for eight hours. Expression was normalised to *CACS* reference transcripts and scaled to the value for mock-treated seedlings within each genotype. Data are means ± SE of *n* = 3 biological replicates. Pairwise significant differences: * *P* < 0.05 ** *P* < 0.01 *** *P* < 0.001; ns, *P* > 0.05 (ANOVA). Source data are provided as a Source Data file.

**Figure 4.**
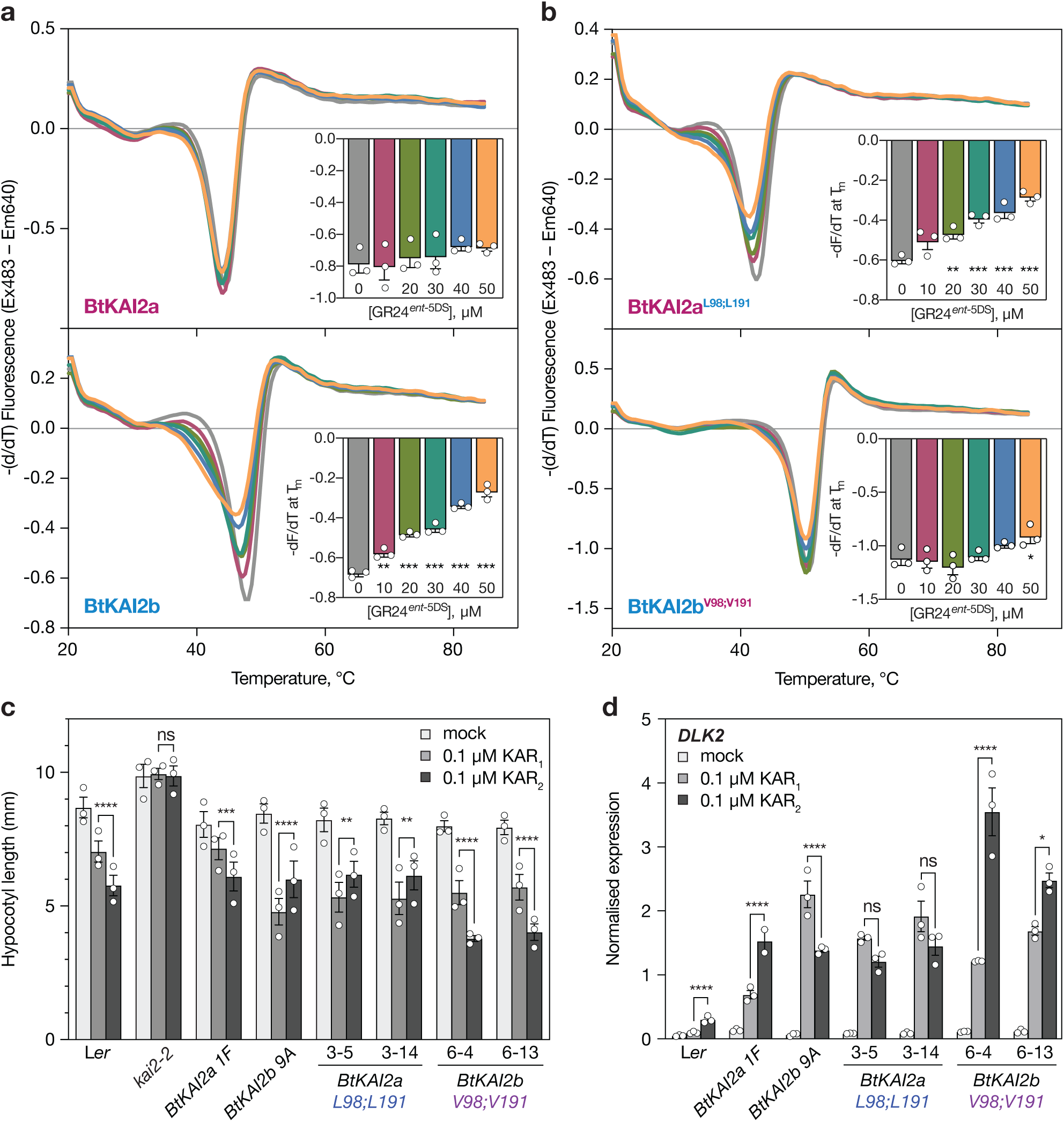
Two residues account for ligand specificity between BtKAI2a and BtKAI2b. **a**, DSF curves of SUMO-BtKAI2a and SUMO-BtKAI2b fusion proteins treated with 0-50 µM GR24*^ent^*^-5DS^, a KAI2-bioactive ligand. Each curve is the average of three sets of reactions, each comprising four technical replicates. Insets plot the minimum value of –(dF/dT) at the melting point of the protein as determined in the absence of ligand (means ± SE, *n* = 3). Significant differences from untreated control: * *P* < 0.05 ** *P* < 0.01 *** *P* < 0.001 (ANOVA). **b**, DSF curves of SUMO-BtKAI2a^L98;L191^ and SUMO-BtKAI2b^V98;V191^ fusion proteins treated with 0-50 µM GR24*^ent^*^-5DS^. **c**, Hypocotyl elongation responses of Arabidopsis expressing GFP-BtKAI2a^L98;L191^ and GFP-BtKAI2b^V98;V191^ fusion proteins and treated with KAR_1_ or KAR_2_. Data are a summary of three experimental replicates performed on separate occasions, each comprising 25-40 seedlings per genotype/treatment combination. Data for each replicate are shown in Supplementary Figure 11. Error bars are SE, n = 3 experimental replicates; each dot corresponds to the mean value derived from each replicate. Asterisks denote significant differences: * P<0.05, ** P<0.01, *** P<0.001, **** P<0.0001 (linear mixed model with experimental replicate as a random effect; specific pairwise comparisons using Tukey’s HSD correction). **d**, Levels of *DLK2* transcripts in the same transgenic lines as above treated with 1 µM KAR_1_, 1 µM KAR_2_, or 0.1% acetone and harvested 8 hours later. Expression was normalised to *CACS* reference transcripts. Data are means ± SE of n = 3 pools of ∼50 seedlings treated in parallel. Asterisks denote significant differences as above (two-way ANOVA; specific pairwise comparisons using Tukey’s HSD correction). Source data are provided as a Source Data file.

We performed transgenic complementation of the Arabidopsis *kai2-2* null mutant by expressing each of the three isoforms as a GFP fusion protein driven by the Arabidopsis *KAI2* promoter and 5′UTR (*KAI2pro*:*GFP-BtKAI2*) to accurately reflect the native expression profile of KAI2 in Arabidopsis. Such fusions of GFP with KAI2 proteins have been used previously to analyse KAI2 activity^19, 40, 41^. Both *BtKAI2a* and *BtKAI2b* complemented the seedling and leaf phenotypes of *kai2-2*, whereas *BtKAI2c* did not (Fig. 2c-d). GFP-BtKAI2a and GFP-BtKAI2b accumulated at consistent levels when detected using both an anti-KAI2 antibody and an anti-GFP antibody (which negated the effects of sequence differences in the anti-KAI2 epitope region; Fig. 2e and Supplementary Fig. 3). However, we could not detect GFP-BtKAI2c protein in three independent transgenic lines with either antibody. We verified the *GFP-BtKAI2c* transcript sequences expressed in the transgenic plant material by RT-PCR and Sanger sequencing (Supplementary Fig. 5). Since the *GFP-BtKAI2c* mRNA is processed faithfully in Arabidopsis, and the apparent absence of protein is most likely a result of posttranslational events. We then tested whether BtKAI2c was generally poorly expressed in plant cells by transient overexpression in tobacco leaves using *Agrobacterium*-mediated infiltration and tobacco mosaic virus-based plant expression vectors^42^. Even with the extremely high levels of expression supported by this system, levels of myc-tagged BtKAI2c protein were substantially lower than equivalently-tagged BtKAI2a and BtKAI2b (Supplementary Fig. 5). This result implies that BtKAI2c protein may be inherently unstable and/or prone to degradation in plant cells.

### BtKAI2c carries a destabilising mutation at position 98

Numerous mutations identified by forward genetics destabilise AtKAI2 and render the protein undetectable in plant extracts^43^. Ten residues in the BtKAI2c sequence are unique to BtKAI2c within the genus *Brassica* (Supplementary Figure 3). We predicted the likely effect of each of these residues upon protein function, relative to the *Brassica* consensus residues at the same positions, using the PROVEAN tool^44^. Among the ten sites, R98 and E129 exceeded the cutoff value of -2.5 and were thus predicted to be highly deleterious to protein function, while D137 narrowly missed this threshold (Supplementary Table 2). Therefore based on this predictive analysis, the native BtKAI2c protein carries at least two and possibly three substitutions that have the potential to render the protein non-functional.

R98 is potentially highly deleterious because it is situated in the active site adjacent to the catalytic serine, whereas E129 and D137 are located on the flexible hinge that links the lid and core domains (Supplementary Fig. 6a). In AtKAI2, the G133E mutation is highly destabilising^20, 41^, suggesting that non-conserved substitutions in the hinge region have the potential to be deleterious as well. To test the effect of these residues on BtKAI2c stability, we constructed three variants that reverted the native BtKAI2c sequence to consensus amino acids within *Brassica* at position 98 and/or in the hinge region: BtKAI2c^R98V^, a quadruple mutant BtKAI2c^E129D;D130V;E133Q;D137E^ (BtKAI2c^QUAD^), and a quintuple mutant that combined all five substitutions (BtKAI2c^QUINT^). Should these amino acids be crucial to protein stability, we expected to restore BtKAI2c expression.

First, we tried expressing the native and mutated BtKAI2c variants in tobacco leaves using *Agrobacterium*-mediated infiltration. The vectors were based on those used to generate the *GFP-BtKAI2c* Arabidopsis transgenics, allowing us to compare expression of the same GFP fusion proteins. As was the case for stably-transformed Arabidopsis, we found that native GFP-BtKAI2c protein was undetectable, as was the BtKAI2c^QUAD^ variant (Supplementary Fig. 6b). However, BtKAI2c^R98V^ and BtKAI2c^QUINT^ were clearly detected. We also examined the effect of the equivalent V96R mutation on GFP-AtKAI2, and found that this mutation completely abolished expression compared to the native GFP-AtKAI2 control (Supplemental Fig. 6b). These results suggest that the R98 residue causes the loss of protein expression in BtKAI2c, and that arginine at this position is not tolerated by KAI2 proteins in general.

Early in our investigation, we had tried without success to express and purify BtKAI2c from *E. coli* for biochemical characterisation. Having restored BtKAI2c expression in plants with the R98V mutation, we next tried to express these proteins in bacteria. We found that SUMO-BtKAI2c R98V was expressed at ∼10-fold higher levels than native SUMO-BtKAI2c in crude lysates, suggesting that the R98V mutation enhanced protein folding and/or stability in bacterial cells. Although it was detectable in crude lysates, SUMO-BtKAI2c was consistently intransigent to recovery following affinity chromatography; in contrast, the R98V and quintuple mutant versions were stably recovered at high levels and purity (Supplemental Figure 6c). From these results, we conclude that native BtKAI2c is non-functional due to protein instability induced by a highly non-conservative substitution at the active site.

### BtKAI2a and BtKAI2b show differential ligand specificity

We further characterised the ligand specificity of *BtKAI2* homologues by performing physiological and molecular assays with the stable transgenic Arabidopsis lines. Primary-dormant Arabidopsis seeds homozygous for the transgenes were tested for germination response (Fig. 3a and Supplementary Fig. 7). None of the transgenic lines completely restored germination levels to those of wild type, perhaps reflecting the need for additional, seed-specific regulatory elements not included in our transgenes, or the importance of genomic context for proper *KAI2* expression in seeds. Nevertheless, germination responses to karrikins were evident in the transgenic seeds: germination of *BtKAI2b* seeds was more responsive to KAR_1_ than KAR_2_ at 1 μM, whereas no significant difference was observed for *BtKAI2a* seeds. As expected, *BtKAI2c* seeds were indistinguishable from the *kai2* control and insensitive to karrikins. We also found that levels of *KUF1* transcripts, which are responsive to karrikins in Arabidopsis seeds (Nelson 2010), were only slightly induced by karrikins in *BtKAI2a* transgenic seeds, but were strongly enhanced by KAR_1_ in *BtKAI2b* seeds (Fig. 3b).

In seedlings, both *BtKAI2a* and *BtKAI2b*, but not *BtKAI2c*, could complement the *kai2* hypocotyl elongation phenotype (Fig 3c). Responses to karrikins in terms of hypocotyl elongation (Fig. 3c) and levels of *DLK2* transcripts (Fig. 3d) broadly agreed with the germination response with respect to karrikin preferences, with *BtKAI2b* transgenics showing a clear preference for KAR_1_ over KAR_2_. However, *BtKAI2a* seedlings showed a preference for KAR_2_ that was not evident in seeds. From these experiments, we conclude that BtKAI2a is similar to AtKAI2 in terms of ligand specificity, whereas BtKAI2b has a clear preference for KAR_1_ over KAR_2_. As *BtKAI2b* is more highly expressed than *BtKAI2a* in *B. tournefortii* seeds and seedlings (Fig. 2b), we conclude that the ligand specificity of BtKAI2b substantially contributes to the enhanced KAR_1_-responsiveness of this species at these stages of the life cycle.

### Positions 98 and 191 account for ligand specificity between BtKAI2a and BtKAI2b

To investigate interactions between BtKAI2 homologues and ligands, we performed differential scanning fluorimetry (DSF) assays on purified recombinant proteins (Supplementary Fig. 8). DSF has been used extensively for inferring the interaction of strigolactone-like compounds with D14- and KAI2-related proteins^29, 41, 45–48^. Racemic GR24 is a widely-used synthetic strigolactone analogue that consists of two enantiomers (Supplementary Fig. 1a), of which GR24*^ent^*^-5DS^ is bioactive via AtKAI2^40, 49^. Catalytically inactive D14 and KAI2 variants do not respond to GR24 in DSF, suggesting that the shift in thermal response results from ligand hydrolysis and a corresponding conformational change in the receptor^29, 41, 46^. In DSF assays, AtKAI2 shows a specific response to >100 µM GR24*^ent^*^-5DS^ but, for unclear reasons, no response to karrikins^41^. Likewise, we found that both BtKAI2a and BtKAI2b were also unresponsive to karrikins in DSF (Supplementary Fig. 9a). Therefore, we used DSF assays with GR24*^ent^*^-5DS^ as a surrogate substrate as a means to analyse differences in ligand interactions between BtKAI2a and BtKAI2b. We used N-terminal 6xHIS-SUMO as a solubility tag to aid expression in *E. coli* and enhance stability. We found that the presence of the tag did not compromise the thermal destabilisation response of BtKAI2b to GR24*^ent^*^-5DS^ (Supplementary Fig. 9b), and therefore we used intact SUMO fusion proteins for all other experiments.

We found that AtKAI2 and BtKAI2a showed similar responses to >100 µM GR24*^ent^*^-5DS^, but the response of BtKAI2b was clear at >25 µM (Supplementary Fig. 9c). We then used a lower range of GR24*^ent^*^-5DS^ concentrations (0–50 µM) to determine the threshold for response, which we defined as a statistically significant reduction in the maximal rate of change in fluorescence at the melting temperature of the protein (T_m_). Although BtKAI2a showed only a weak and non-significant response at 40 and 50 µM GR24*^ent^*^-5DS^, BtKAI2b responded significantly at 10 µM and above (Fig. 4a). These results suggest that BtKAI2b is more sensitive than BtKAI2a to GR24*^ent^*^-5DS^, and that BtKAI2a is most like AtKAI2 in this respect.

BtKAI2a and BtKAI2b differ in primary amino acid sequence at just 14 positions (Supplementary Fig. 3). We postulated that differences in ligand specificity might be determined by amino acids in the vicinity of the ligand binding pocket. Protein structural homology models revealed only two residues that differ in this region: V98 and V191 in BtKAI2a, and L98 and L191 in BtKAI2b; the corresponding residues in AtKAI2 and AtD14 are valines (Supplementary Fig. 8). Residue 98 is immediately adjacent to the catalytic serine at the base of the pocket. Residue 191 is located internally on αT4 of the lid domain which, in AtD14, is associated with a major rearrangement of protein structure upon ligand binding that reduces the size of the pocket^31^. Homology modelling suggests only subtle differences in the size and shape of the primary ligand binding pocket of BtKAI2a and BtKAI2b (Supplementary Fig. 8). To determine if these residues are pertinent to ligand specificity, we replaced the two valine residues of BtKAI2a with leucine residues, generating the variant BtKAI2a^L98;L191^, and *vice-versa* for BtKAI2b, generating the variant BtKAI2b^V98;V191^ (Supplementary Fig. 8). In DSF assays, we found that exchanging the two residues was sufficient to switch the original responses, such that BtKAI2a^L98;L191^ responded sensitively to GR24*^ent^*^-5DS^, but BtKAI2b^V98;V191^ did not (Fig. 4b). We also assessed the function of the stable BtKAI2c^R98V^ and BtKAI2c^QUINT^ variants by DSF against GR24*^ent^*^-5DS^ ligand, and found that they were only weakly destabilised by this ligand, when compared to the highly sensitive BtKAI2b (Supplemental Fig. 6e). Overall, these results indicate that BtKAI2a and BtKAI2b have distinct response profiles, albeit to a synthetic ligand, and that residues 98 and 191 contribute to this difference.

We reasoned that the most robust and informative means to assess the effect of residues 98 and 191 upon the response to karrikins was to test their function directly *in planta*. We expressed BtKAI2a^L98;L191^ and BtKAI2b^V98;V191^ as GFP fusion proteins in the Arabidopsis *kai2-2* null mutant background driven by the Arabidopsis *KAI2* promoter (*KAI2pro:GFP-BtKAI2a^L98;L191^* and *KAI2pro:GFP-BtKAI2b^V98;V191^*), and selected two independent homozygous transgenic lines for each, on the basis of protein expression level and segregation ratio (Supplementary Fig. 10). Using two different assays, we found that substitutions between BtKAI2a and BtKAI2b at positions 98 and 191 also reversed karrikin preference in Arabidopsis seedlings both in terms of hypocotyl elongation and *DLK2* transcript levels (Fig. 4c, d; Supplementary Fig. 11). Most prominently, the clear preference of BtKAI2b for KAR_1_ was unambiguously reversed to a preference for KAR_2_ in *BtKAI2b^V98;V191^* transgenics, effectively recapitulating the response of the native BtKAI2a protein. We also examined the responses of seeds in these transgenic lines, and found that *BtKAI2a^L98;L191^* seeds germinated with a clear preference for KAR_1_, whereas *BtKAI2b^V98;V191^* seeds showed no clear preference for either karrikin (Supplementary Fig. 12). Notably, this germination response pattern is the opposite to that observed for seeds expressing the native BtKAI2 proteins, with BtKAI2a exhibiting no preference for either karrikin and BtKAI2b a preference for KAR_1_ (Fig. 3a, Supplementary Fig. 7). Taken together, these results demonstrate that the residues at positions 98 and 191 largely determine differences in karrikin specificity between BtKAI2a and BtKAI2b, at least in the context of Arabidopsis seedlings.

### Taxonomic variability in KAI2 sequences suggests functional co-dependency between residues 96 and 189

We wished to gain further insight into the functional importance of diversity at positions 96 and 189 of KAI2 proteins (based on AtKAI2 notation, equivalent to positions 98 and 191 in BtKAI2a and BtKAI2b). We analysed 476 distinct KAI2 homologues from 441 angiosperm species (spanning eudicots, monocots and magnoliids, but excluding *B. tournefortii*) using data collected from the 1000 Plant Transcriptomes project^50^. We found that at positions 96 and 189, valine was by far the most common amino acid (93.1% and 93.5% of sequences, respectively; Figure 5a and Supplemental Table 3). L96 was observed in 6.1% of sequences, while L189 was even rarer (2.1%). The L96;V189 combination was observed in 14 sequences (2.9%), whereas V96;L189 was observed in only one homologue from *Ternstroemia gymanthera* (Pentaphylacaceae; 0.2%). Notably, this species also expressed a second KAI2 homologue with the canonical V96;V189 combination. If the frequencies of valine and leucine at positions 96 and 189 were independent, the L96;L189 combination would be expected to occur less than once among 476 sequences (0.13%). However, the coincidence of L96 and L189 was observed in nine sequences (1.9%), representing a significant 15-fold enrichment for the L96;L189 pair over that expected by chance alone (χ^2^ = 115.6; P<0.0001).

**Figure 5.**
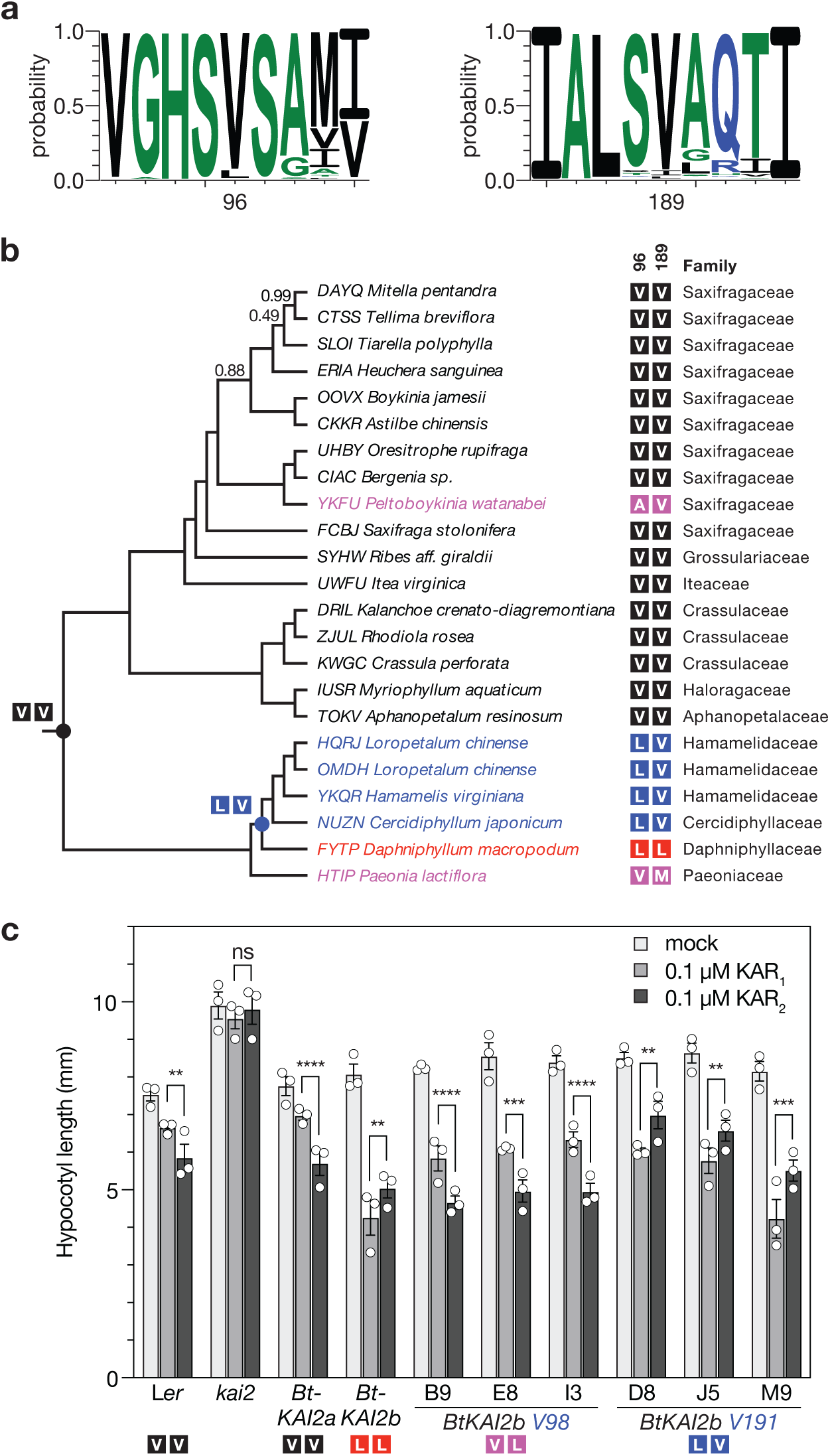
Diversity at positions 96 and 189 in KAI2 homologues across angiosperms. **a**, Sequence logos generated from 476 aligned KAI2 homologues sampled from eudicots, monocots and magnoliids and centred on position 96 (left) and 189 (right). b, Species tree of the Saxifragales based upon 410 single-copy nuclear genes estimated using the ASTRAL gene tree method^50^. Branch support values are shown only where less than 1. The four-letter taxon prefix corresponds to the 1000 Plant Transcriptomes project Sample ID. Taxa are highlighted by colour according to identity of amino acids of KAI2 orthologues at positions 96 and 189 respectively: black, VV; blue, LV; red, LL; magenta, other. Postulated ancestral identities at specific nodes are indicated with filled circles. c, Hypocotyl elongation responses to karrikins in three independent transgenic lines homozygous for *GFP-BtKAI2a^L98;V191^* and three lines for *GFP-BtKAI2b^V98;L191^*. Data shown are a summary of three experimental replicates performed on separate occasions, each comprising ∼20 seedlings per genotype/treatment combination. Data from each replicate are presented in Supplementary Figure 13. Error bars are SE, n = 3 experimental replicates; each dot corresponds to the mean value derived from each replicate. Asterisks denote significant differences: * P<0.05, ** P<0.01, *** P<0.001, **** P<0.0001 (linear mixed model with experimental replicate as a random effect; specific pairwise comparisons using Tukey’s HSD correction). Source data are provided as a Source Data file.

Among the nine newly-identified KAI2^L96;L189^ sequences, eight were restricted to eudicots, distributed evenly between both the superrosids (Brassicales, Fabales and Saxifragales; four species) and superasterids (Caryophyllales; four species). The ninth sequence, co-expressed with another transcript encoding KAI2^V96;V189^, was identified from the basal dicot *Hakea drupacea* (syn. *Hakea suaveolens*; Proteaceae). Strikingly, this species is native to southwestern Australia, but is recognised as an invasive weed in South Africa that relies upon fire to liberate seeds from the canopy^51^. The L96;L189 combination therefore seems to have arisen independently on multiple occasions. Furthermore, within the order Saxifragales, *Daphniphyllum macropodum* (KAI2^L96;L189^) belongs to a clade of species that all express KAI2^L96;V189^ homologues (Figure 5b). Although this clade is only represented by five sequences and four species, this distribution pattern suggests that a V96L mutation occurred first in the common ancestor of the Hamamelidaceae, Cercidiphyllaceae and Daphyniphyllaceae, followed by V189L in the lineage leading to *D. macropodum*. Overall, this analysis suggests that positions 96 and 189, while not in direct contact with each other in the protein tertiary structure (Supplemental Fig. 8), likely exhibit functional interdependence. Valine is most probably the ancestral state at both positions, but leucine at position 96 is also tolerated. While rare, leucine at position 189 is generally found in combination with leucine at position 96, and tentatively, a V96L mutation might potentiate a subsequent V189L mutation.

The above analysis suggested that a V96L substitution might be sufficient to confer altered substrate specificity upon KAI2 proteins. To test this hypothesis, we generated transgenic Arabidopsis seedlings expressing GFP-BtKAI2b proteins with just a single amino acid substitution from leucine to valine at either position 98 or 191 (i.e. *KAI2pro:GFP-BtKAI2b^V98^* or *KAI2pro:GFP-BtKAI2b^V191^*), and examined their response to KAR_1_ and KAR_2_ in hypocotyl elongation assays. We found that the L98V substitution was sufficient to invert the karrikin preference of GFP-BtKAI2b in three independent homozygous transgenic lines (Figure 5c; Supplemental Figure 13). Although this result does not rule out an additional contribution towards ligand specificity from L191, it is consistent with the higher frequency of L96 compared to L189 observed among the angiosperms, and suggests that substitution at position 96 may have been selected relatively frequently in the evolution of KAI2 function.

## DISCUSSION

The plant α/β-hydrolases KAI2 and D14 are characterised by responsiveness towards butenolide compounds, including endogenous strigolactones and strigolactone-related compounds, abiotic karrikins derived from burnt vegetation, and synthetic strigolactone analogues with a wide array of functional groups. Direct evidence for a receptor-ligand relationship between karrikins and KAI2 homologues stems from crystallography and *in vitro* binding assays. Two crystal structures of KAR-responsive KAI2 proteins from Arabidopsis and the parasitic plant *Striga hermonthica* reveal largely similar overall protein structure, but surprisingly are non-congruent with respect to KAR_1_ binding position and orientation^52, 53^. The affinity of AtKAI2 for KAR_1_ is imprecisely defined, with estimates of dissociation coefficients (Kd) ranging from 4.6 µM^54^, to 26 µM^55^, to 148 µM^56^ using isothermal calorimetry, and 9 µM using fluorescence microdialysis^52^. While variability in affinity estimates can be explained in part by different experimental conditions and techniques, it should also be considered that, depending on the homologue under examination, KAR_1_ may not be the optimal ligand. Furthermore, binding of a hydrophobic compound to a hydrophobic pocket does not necessarily equate to ligand recognition or receptor activation. Our data, which are derived from clear and distinct biological responses, provide strong evidence that KAI2 is sufficient to determine ligand specificity, which in turn strengthens the case that KAI2 is the receptor by which karrikins are perceived.

The predominance of valine and leucine at position 96 (98 in BtKAI2 proteins), observed in 99.2% of our sample of 476 KAI2 sequences, suggests that this site is critical to protein function, which makes sense given its immediate proximity to the catalytic serine. This position not only contributes to ligand preference, but also to protein stability, given the negative effects of R98 in BtKAI2c. Compared with valine and leucine, arginine is relatively bulky with a charged side chain; indeed, in the few KAI2 sequences in which position 96 is neither leucine nor valine, the residue is nevertheless small and hydrophobic (either methionine or alanine; Supplementary Table 3). Interestingly, substitutions in AtKAI2 at or near the catalytic site, such as S95F, S119F and G245D, are not tolerated and render the protein unstable in plants^43^. The likely signalling mechanism for KAI2 – by analogy of that for D14 – involves ligand binding and/or hydrolysis, which triggers conformational changes in the lid domain and formation of a protein complex with MAX2 and SMAX1/SMXL2 protein partners^30, 31, 57^. As a result of signalling, KAI2 itself is degraded^40, 43^. It is possible that KAI2 is an inherently metastable protein that is prone to conformational change and subsequent degradation by necessity; perhaps this explains why it is so susceptible to instability through mutation.

Ligand specificity is a key contributor to the functional distinction between karrikin and strigolactone receptors, and elucidating the molecular mechanisms behind ligand specificity is a significant research challenge. In certain root-parasitic weeds in the Orobanchaceae, substitutions of bulky residues found in AtKAI2 with smaller hydrophobic amino acids that increase the ligand-binding pocket size have likely improved the affinity for host-derived strigolactone ligands, as opposed to the smaller-sized karrikin-type ligands^58–60^. Moreover, lid-loop residues that affect the rigidity and size of the ligand entry tunnel determine the ligand selectivity between KAR_1_ and *ent*-5-deoxystrigol (a strigolactone analogue with non-natural stereochemistry) among eleven KAI2 homologues in *Physcomitrella patens*^61^. Karrikin and strigolactone compounds also show chemical diversity within themselves, yet ligand discrimination among KAI2 and D14 receptors is not well characterised. Our data demonstrate that, in the case of BtKAI2 proteins, the two KAI2 homologues respond differently to subtly different KAR analogues, and that subtle changes in pocket residues likely account for preferences between these ligands. Position 96 appears to play a primary role in determining ligand specificity, while position 189 may improve ligand sensitivity, protein stability or protein conformational dynamics following a prior mutation at position 96. Although the identified residues L98 and L191 are not predicted to change dramatically the pocket size of BtKAI2b in comparison to BtKAI2a, these residues would make the BtKAI2b pocket more hydrophobic, which is consistent with KAR_1_ being more hydrophobic than KAR_2_^4^. Therefore, ligand specificity between highly similar chemical analogues can be achieved through fine-tuning of pocket hydrophobicity. Similarly subtle changes have been reported for the rice gibberellin receptor GID1: changing Ile133 to Leu or Val increases the affinity for GA34 relative to the less polar GA4, which lacks just one hydroxyl group^62^.

It is possible that what we learn about ligand specificity in KAI2 proteins may also be informative about strigolactone perception by D14. Different D14 proteins may show ligand specificity towards diverse natural strigolactones, resulting in part from multiplicity of biosynthetic enzymes, such as the cytochrome P450 enzymes in the MAX1 family^45, 63, 64^ and supplementary enzymes such as LATERAL BRANCHING OXIDOREDUCTASE^65^. Although the functional reasons underlying such strigolactone diversity are still unclear, it is possible that variation among D14 homologues yields varying affinities for different strigolactone analogues^66^. As an increasingly refined picture emerges of the features that determine ligand specificity for KAI2 and D14 receptors, we envision the rational design of synthetic receptor proteins with desirable ligand specificity in the future.

Gene duplication is a common feature in plant evolutionary histories as an initial step towards neofunctionalisation^67^. In obligate parasitic weeds, duplication and diversification of *KAI2* genes (also referred to as *HTL* genes) have shifted ligand specificity towards strigolactones^58, 59^. Karrikins, as abiotic molecules with limited natural occurrence, are unlikely to be the natural ligand for most KAI2 proteins, which are found throughout land plants. Instead, the evolutionary maintenance of a highly conserved receptor probably reflects the core KAI2 function of perceiving an endogenous ligand (“KL”) that regulates plant development^34–36^. This core function is presumably retained after gene duplication events through original, non-divergent copies that are under purifying selection^58, 59^. KAI2 diversity may also reflect differing affinity for KL variants in different species and at different life stages. Our results are consistent with a scenario in which both BtKAI2a and BtKAI2b have retained the core KAI2 function of perceiving KL to support plant development, because both copies complement the Arabidopsis *kai2* phenotype, especially at the seedling and later stages. However, BtKAI2b has also acquired mutations that alter its ligand specificity, which in turn enhance sensitivity to the archetypal karrikin released first discovered in smoke. The *BtKAI2b* gene is also more highly expressed than *BtKAI2a* in seeds and seedlings, consistent with an adaptation to post-fire seedling establishment. This diversity among KAI2 proteins may provide *B. tournefortii* with a selective advantage in fire-prone environments, contributing to its invasive nature.

We have shown evidence for the independent evolution of valine-to-leucine substitutions in KAI2 homologues throughout the dicots, albeit at low frequency, suggesting that these changes may have adaptive significance in various ecological contexts. It will be of interest to assess the function of such homologues in other species, to validate and extend the specific conclusions we have made here. Recent findings indicate that KAI2 is an integrator that also modulates germination in response to other abiotic environmental signals, including temperature and salinity^68^. As germination is a critical life stage for seed plants, strategies for environmental conservation, restoration and weed control will benefit from specific knowledge of KAI2 sequence diversity and expression profiles.

## METHODS

### Chemical synthesis

Karrikins (KAR_1_ and KAR_2_), ^13^[C]_5_-labelled karrikins and GR24 enantiomers (GR24^5DS^ and GR24*^ent^*^-5DS^) were prepared as previously described^69, 70^.

### Plant material

Arabidopsis *kai2-2* (L*er*) and *Atd14-1* (Col-0) mutants were previously described^20^. Arabidopsis plants were grown on a 6:1:1 mixture of peat-based compost (Seedling Substrate Plus; Bord Na Mona, Newbridge, Ireland), vermiculite and perlite, respectively. Light was provided by wide-spectrum white LED panels emitting 120-150 µmol photons.m^-2^.s^-1^ with a 16 h light/8 h dark photoperiod, a 22 °C light/16 °C dark temperature cycle, and constant 60% relative humidity. The seeds of *Brassica tournefortii* used in this work were collected in November and December 2009 from two sites in Western Australia (Kings Park in Perth, and Merridin)^71^. *Brassica tournefortii* seeds were dried to 15% relative humidity for one month prior to storage in air-tight bags at –20 °C.

### Seed germination assays

Seed germination assays using Arabidopsis were performed on Phytagel as described previously^49^. *Brassica tournefortii* seeds were sowed in triplicates (35-70 seeds each) on glass microfibre filter paper (Grade 393; Filtech, NSW Australia) held in 9-cm petri dishes and supplemented with mock or karrikin treatments (3 mL of aqueous treatment solution per petri dish). Treatments were prepared by diluting acetone (mock) or karrikin stocks (dissolved in acetone) 1:1000 with ultrapure water. Because germination of *B. tournefortii* is inhibited by light^9^, the seeds were imbibed in the dark at 22 °C. Numbers of germinated seeds were counted each day until the germination percentages remained unchanged.

### Hypocotyl elongation assays

Arabidopsis hypocotyl elongation assays were performed under red light as described previously^20^. For *B. tournefortii*, assays were performed with the following modifications to the Arabidopsis protocol: *B. tournefortii* seeds were sowed in triplicate on glass microfibre filter paper (Filtech) held in petri-dishes supplemented with mock or karrikin treatments. The seeds were imbibed in the dark for 22 h at 24 °C before exposing to continuous red light (5 µmol photons m^-2^ s^-1^) for 4 days.

### Transcript analysis

*Brassica tournefortii* seeds were imbibed on glass fibre filters in the dark at 22 °C and treated with karrikins using the same procedure as described under seed germination assays, above. For transcript quantification in *B. tournefortii* seeds, samples were taken at the indicated time points, briefly blot-dried on paper towel, and frozen in liquid nitrogen. For *B. tournefortii* seedlings, two methods were used. For data shown in Fig. 1e, seeds were sowed and seedlings grown in triplicate under identical conditions to those described for hypocotyl elongation assays. For data shown in Fig. 2b, seeds were first imbibed for 24 h in the dark, and then transferred to the growth room (∼120-150 μmol photons m^-2^ s^-2^ white light, 16 h light/8 h dark, 22 °C light/16 °C dark) for three days. A sample of approximately 20 seedlings was then transferred to a 250 mL conical flask containing 50 mL sterile ultrapure water supplemented with 1 µM KAR_1_, or an equivalent volume of acetone (0.1% v/v), with three replicate samples per treatment. The flasks were then shaken for 24 h before seedlings were harvested.

Arabidopsis seedlings were grown on 0.5× MS agar under growth room conditions (wide-spectrum white LED panels emitting 120-150 µmol photons.m^-2^.s^-1^ with a 16 h light/8 h dark photoperiod and a 22 °C light/16 °C dark temperature cycle) for seven days. On the seventh day, seedlings were transferred to 3 mL liquid 0.5× MS medium in 12-well culture plates (CORNING Costar 3513) and shaken at 70 rpm for a further 22 h under the same growth room conditions. The medium was then removed by pipette and replaced with fresh medium containing relevant compounds or an equivalent volume (0.1% v/v) of acetone. After a further incubation period with shaking (8 h), the seedlings were harvested, blotted dry, and frozen in liquid nitrogen.

RNA extraction, DNase treatment, cDNA synthesis and quantitative PCR were conducted as previously described^72^. All oligonucleotides are listed in the Supplementary Table 4.

### Cloning and mutagenesis of *Brassica tournefortii KAI2* and *D14* homologues

Full-length *BtKAI2* coding sequences (and unique 3′UTR sequences) were amplified from *B. tournefortii* cDNA with Gateway-compatible *attB* sites using the universal forward primer BtKAI2_universal_F and homologue-specific reverse primers RACE_R (BtKAI2a), Contig1_R (BtKAI2b) and Contig5_R (BtKAI2c), before cloning into pDONR207 (Life Technologies). The pDONR207 clones were confirmed by Sanger sequencing and recombined with pKAI2pro-GFP-GW^41^ to generate the binary plant expression plasmids pKAI2pro-GFP-BtKAI2a, pKAI2pro-GFP-BtKAI2b and pKAI2pro-GFP-BtKAI2c. For transient overexpression in tobacco (Supplementary Figure 5d), BtKAI2 coding sequences were transferred via Gateway-mediated recombination into pSKI106, which drives very high expression of coding sequences placed immediately downstream of a full-length cDNA clone of the tobacco mosaic virus U1 strain, all under control of the CaMV *35S* promoter in a standard T-DNA binary vector^73^. pSKI106 also encodes an N-terminal 3× *c*-myc tag. For transient expression in tobacco using standard binary vectors (Supplementary Figure 6b), the *BtKAI2* coding sequences were transferred into pMDC43, which is identical to pKAI2pro-GFP-GW except that the *AtKAI2* promoter & 5′UTR sequences are replaced with a double CaMV *35S* promoter^74^. The *BtD14a* coding sequence was amplified from cDNA using oligonucleotides BtD14_F and BtD14_R, cloned into pDONR207 as above, and transferred into pD14pro-GW^41^.

The full-length *BtKAI2* coding sequences (excluding 3′UTRs) were amplified from the pDONR207 clones and reconstituted with the pE-SUMO vector by Gibson Assembly to generate the heterologous expression plasmids pE-SUMO-BtKAI2a, -BtKAI2b and -BtKAI2c. Site-directed mutagenesis generated pE-SUMO-BtKAI2a^L98;L191^, pE-SUMO-BtKAI2b^V98;V191^, and the BtKAI2c variants. For expression of mutated versions of BtKAI2a, BtKAI2b, BtKAI2c and AtKAI2 in tobacco and Arabidopsis, site-directed mutagenesis was performed on pDONR207 clones prior to recombination with pKAI2pro-GFP-GW or pMDC43. In the case of the double substitutions in BtKAI2a and BtKAI2b, both targeted residues were mutated simultaneously in one PCR product, while the remainder of the plasmid was amplified in a second PCR product. These double mutated plasmids were reconstituted by Gibson assembly. For the single mutants, the entire plasmid was amplified using a single pair of mutagenesis primers, and the PCR product was self-ligated. The BtKAI2c^QUINT^ variant was generated by two successive rounds of mutagenesis. Coding regions were confirmed by Sanger sequencing.

### Plant transformation

Homozygous *kai2-2* plants were transformed by floral dip. Primary transgenic seedlings were selected on sterile 0.5× MS medium supplemented with 20 µg/mL hygromycin B. T2 lines exhibiting a 3:1 ratio of hygromycin resistant-to-sensitive seedlings were propagated further to identify homozygous lines in the T3 generation. Experiments were performed from the T3 generation onwards.

For transient expression in tobacco, *Agrobacterium* (GV3101) carrying pSKI106 or pMDC43 variants was grown in LB medium (25 mL) supplemented with antibiotics and 20 µM acetosyringone until OD600 reached 1.0. The bacteria were then harvested by centrifugation (15 min, 5000 × *g*) and resuspended in 10 mM MgCl2, 10 mM MES (pH 5.6) and 100 µM acetosyringone. The optical density was adjusted to 0.4, and the suspension was left standing at 22 °C overnight (approximately 14 h). Leaves of three-week-old *Nicotiana benthamiana* were then infiltrated with a 5-mL syringe, through the abaxial leaf surface. After four days, the leaves were collected and frozen in liquid nitrogen.

### Karrikin uptake measurements

Fifteen samples of seeds were sowed for each karrikin treatment (five time points, each in triplicate). In each sample, approximately 40 mg of *Brassica tournefortii* seeds were imbibed in 3 mL ultrapure water for 24 h in 5-mL tubes. After centrifugation (2 min at 3220 × *g*), excess water was removed by pipette and the volume of residual water (mostly absorbed into the seeds) was calculated by weighing the seeds before and after imbibition. Fresh ultrapure water was added to the fully-imbibed seeds to reach a total volume of 980 μL. Then 20 μL of 100 μM KAR_1_ or KAR_2_ was added to a final concentration of 2 μM. The seeds were imbibed at 22 °C in darkness. At the indicated time point (0, 2, 4, 8 or 24 h post-treatment), 500 μL of the imbibition solution was removed and combined with 100 ng of either ^13^[C]_5_-KAR_1_ or ^13^[C]_5_-KAR_2_ (100 μL at 1 μg/mL) as an internal standard for quantification purposes. The sample was then extracted once with ethyl acetate (500 μL), and 1 μL of this organic layer was analysed using GC-MS in selective ion monitoring (SIM) mode as previously described^75^.

The amount of KAR_1_ in each sample was calculated by the formula 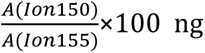 and converted to moles, where *A(Ion150)* indicates the peak area of the ion 150 (KAR_1_ to be measured), *A(Ion151)* indicates the peak area of the ion 151 (^13^[C]_5_-KAR_1_), and 100 ng is the amount of ^13^[C]_5_-KAR_1_ spiked in before the ethyl acetate extraction. Similarly, the amount of KAR_2_ in each sample was calculated by the formula 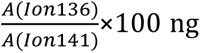 and converted to moles. The uptake percentage adjusted to 40 mg of *B. tournefortii* seeds was calculated by the formula: 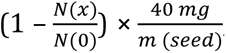, where N(0) indicates moles of karrikins at time 0, *N*(*x*) indicates moles of karrikins at time point *x*, and *m*(*seed*) indicates the dry weight (mg) of seeds tested in each replicate. For Arabidopsis seeds, the procedure was scaled down for a smaller mass of seeds (20 mg).

### Transcriptome assembly and analysis

Twenty milligrams of dry *B. tournefortii* seeds were imbibed in water for 24 h and incubated at 22 °C in the dark. The seeds were collected by centrifugation, blotted dry, and frozen in liquid nitrogen. A separate sample of seeds was sown on glass filter paper, imbibed for 24 h as above, and then incubated for 96 h under continuous red light (20 µmol m^-2^ s^-1^) with a 22 °C (16 h)/16 °C (8h) temperature cycle. A single sample of seedlings (50 mg fresh weight) was harvested and frozen in liquid nitrogen. Total RNA was extracted from both seed and seedling samples using the Spectrum Plant RNA kit (Sigma-Aldrich), including an on-column DNase step. PolyA^+^ mRNA was purified using oligo(dT) magnetic beads (Illumina), and cDNA libraries for sequencing were generated as described^76^. Sequencing was performed on the Illumina HiSeq 2000 platform at the Beijing Genomic Institute, Shenzhen, China. Raw reads were filtered to remove adapters and low-quality reads (those with >5% unknown nucleotides, or those in which greater than 20% of base calls had quality scores ≤10). After filtering, both libraries generated reads with >99.9% of nucleotides attaining Q20 quality score. Transcriptome *de novo* assembly was performed with Trinity^77^. For each library, contigs were assembled into Unigenes; Unigenes from both libraries were then combined, yielding a total of 45,553 predicted coding region sequences with a mean length of 1011 nt. The combined Unigenes were then interrogated for homology to *AtKAI2*, *AtD14*, *AtDLK2*, *AtSTH7* and *AtCACS* using BLASTn searches.

### Phylogenetic analysis and protein sequence analysis

*KAI2* and *D14* homologues in *Brassica* species were identified from BLAST searches using Arabidopsis coding sequences as a query. Additional sequences were sampled from an existing phylogenetic analysis^33^. Multiple sequence alignments were performed using MAFFT plugin implemented in Geneious R10 (Biomatters). The alignment was trimmed slightly at the 5′ end to remove non-aligned regions of monocot D14 sequences. Maximum likelihood phylogenies were generated using PHYML (GTR +G +I substitution model, combination of NNI and SPR search, and 100 bootstraps). The choice of substitution model was guided by Smart Model Selection in PhyML^78^ (http://www.atgc-montpellier.fr/sms). A list of all sequences, and their sources, is provided in Supplementary Table 1.

For analysis of BtKAI2c using PROVEAN, all ten positions that were unique to BtKAI2c when compared to AtKAI2 and other *Brassica* sequences (Supplementary Fig. 3) were manually edited to the consensus identity at each position. This was the “reverted” sequence against which each of the native BtKAI2c amino acid substitutions were tested using the PROVEAN server (http://provean.jcvi.org) using the default cut-off score of -2.5. The PROVEAN output is presented in Supplementary Table 2.

KAI2 homologues from The 1000 Plant Transcriptomes Project^50^ were identified by TBLASTN searches using *Arabidopsis thaliana* KAI2 protein sequence as a query against “onekp database v5”, which comprises 1328 individual samples (https://db.cngbdb.org). The search was configured to return a maximum of 500 target sequences with Expect values <0.01, using a word size of 6 and the BLOSUM62 scoring matrix. Nucleotide sequences were collated, and open reading frames (ORFs) >700 nucleotides were predicted in Geneious software. A single ORF was selected for each sequence; those with multiple ORFs were screened manually for the one with homology to AtKAI2. The ORFs were translated and aligned using the MAFFT multiple alignment tool. A few misaligned or mis-translated sequences were removed from the alignment manually, resulting in a dataset of 476 unique sequences. Extracted parts of this alignment centred on positions 96 and 189 were used to generate WebLogos (http://weblogo.threeplusone.com79). A list of identified KAI2 homologues and their positional analysis are presented in Supplementary Table 3.

### Protein homology modelling

KAI2 structures were modelled using the SWISS-MODEL server (https://swissmodel.expasy.org) using the alignment mode^80^ and the *Arabidopsis thaliana* KAI2 structure 3w06 as a template^54^. Figures of protein structure and homology models were generated using PyMOL v1.3 (Schrödinger LLC). Cavity surfaces were visualised using the “cavities & pockets (culled)” setting in PyMOL and a cavity detection cut-off value of four solvent radii. Cavity volumes were calculated using the CASTp server v3.0^81^ (http://sts.bioe.uic.edu/castp) with a probe radius of 1.4Å. Values indicate Connolly’s solvent-excluded volumes. Cavities were inspected visually using Chimera v1.12 (https://www.cgl.ucsf.edu/chimera/). For both BtKAI2a and BtKAI2b models, CASTp erroneously included surface residues near the primary pocket entrance in the calculation of the pocket volumes. This issue was resolved by the artificial placement of a free alanine residue adjacent to the cavity entrance, as described previously^82^.

### Protein expression and purification

BtKAI2 proteins were generated as N-terminal 6×HIS-SUMO fusion proteins. All proteins were expressed in BL21 Rosetta DE3 pLysS cells (Novagen) and purified using IMAC as described in detail previously^41^.

### Differential scanning fluorimetry

DSF was performed in 384-well format and thermal shifts were quantified as described previously^41^. Reactions (10 µL) contained 20 µM protein, 20 mM HEPES pH 7.5, 150 mM NaCl, 1.25% (v/v) glycerol, 5× SYPRO Tangerine dye (Molecular Probes) and varying concentrations of ligand that resulted in a final concentration of 5% (v/v) acetone.

### Statistical analysis

Data were analysed using one- or two-way ANOVA (α = 0.05, with Tukey’s multiple comparisons test). For Figure 5a, in which data from three experimental replicates were combined, data were analysed using a mixed effects model with experimental replicate as a random effect, and genotype and treatment as fixed effects. Prior to ANOVA, germination data were arcsine-transformed, and gene expression data were log-transformed. Tests were implemented in GraphPad Prism version 7.0 or 8.0 (GraphPad Software, graphpad.com).

### DATA AVAILABILITY

All relevant data and materials are available from the authors upon reasonable request. Sequence data are available at NCBI Genbank under the following accessions: *BtKAI2a*, MG783328; *BtKAI2b*, MG783329; *BtKAI2c*, MG783330; *BtD14a*, MG783331; *BtD14b*, MG783332; *BtDLK2*, MG783333; *BtSTH7*, MK756121; *BtCACS*, MK756122. Raw RNA sequence data from *Brassica tournefortii* seed and seedlings are available in the NCBI SRA database under accession SRP128835. The original data underlying the following figures are provided as a Source Data file: Figs 1a,b,d,e; 2b,e; 3a–d; 4a–d; 5a–c; Supplementary Figs 1b–f; 2; 3; 4c; 5b–d; 6b,c,e; 7; 8a–e; 9a–c; 10a; 11; 12; 13. A reporting summary for this Article is available as a Supplementary Information File.

## ACKNOWLEDGEMENTS

This work was supported by funding from the Australian Research Council (DP130103646 to SMS and GRF; DP140104567 to SMS; FT150100162 to MTW, DP160102888 to GRF and MTW). Y.K.S was recipient of a Research Training Program Scholarship from the Australian Government. We thank Dr Rowena Long for the original provision of *B. tournefortii* seeds, Dr Rohan Bythell-Douglas for advice on homology modelling, and Mr Yongjie Meng with assistance with RNA extractions. We are grateful to Dr Kevin Rozwadowski (Agriculture and Agri-Food Canada) for the provision of pSKI106.

## CONTRIBUTIONS

Y.K.S., S.M.S., C.S.B., G.R.F. and M.T.W. conceived and designed the research. Y.K.S. and M.T.W. performed the majority of experiments with assistance from J.Y. (hypocotyl elongation assays and screening of transgenic lines), A.S. (chemical synthesis and seed germination assays), K.T.M. and S.F.D. (protein expression and purification). Y.K.S. and M.T.W. analysed the data. Y.K.S., S.M.S. and M.T.W. wrote the manuscript.

## ADDITIONAL INFORMATION

**Supplementary Figure 1** Germination of *Brassica tournefortii* seed treated with karrikins

**Supplementary Figure 2** Extended phylogeny of eu-KAI2 and D14 proteins in angiosperms

**Supplementary Figure 3** Alignment of Brassica KAI2 sequences

**Supplementary Figure 4** BtD14a is functionally homologous to AtD14

**Supplementary Figure 5** The *GFP-BtKAI2c* transgene is faithfully transcribed in Arabidopsis

**Supplementary Figure 6** The R98V mutation restores stability of BtKAI2c

**Supplementary Figure 7** Germination profiles of transgenic Arabidopsis seeds expressing native BtKAI2 homologues

**Supplementary Figure 8** BtKAI2 homology models and SDS-PAGE of SUMO-BtKAI2 fusion proteins used for DSF

**Supplementary Figure 9** BtKAI2a and BtKAI2b do not respond to karrikins in DSF assays

**Supplementary Figure 10** Stable transgenic expression of BtKAI2 valine–leucine double exchange proteins in Arabidopsis

**Supplementary Figure 11** Three experimental replicates of hypocotyl elongation assays with BtKAI2 Arabidopsis transgenics (double exchange of residues 98 and 191)

**Supplementary Figure 12** Germination profiles of transgenic Arabidopsis seed expressing BtKAI2 homologues

**Supplementary Figure 13** Three experimental replicates of hypocotyl elongation assays with BtKAI2b transgenics (individual exchange of residues 98 and 191)

**Supplementary Table 1** List of sequences identified in this study

**Supplementary Table 2** Analysis of BtKAI2c sequence using PROVEAN

**Supplementary Table 3** Analysis of KAI2 homologues identified from The 1000 Plant Transcriptomes project

**Supplementary Table 4** List of oligonucleotides

**Supplementary References**

**Reporting Summary**

**Source Data file**

## SUPPLEMENTARY FIGURE LEGENDS

**Supplementary Figure 1.**
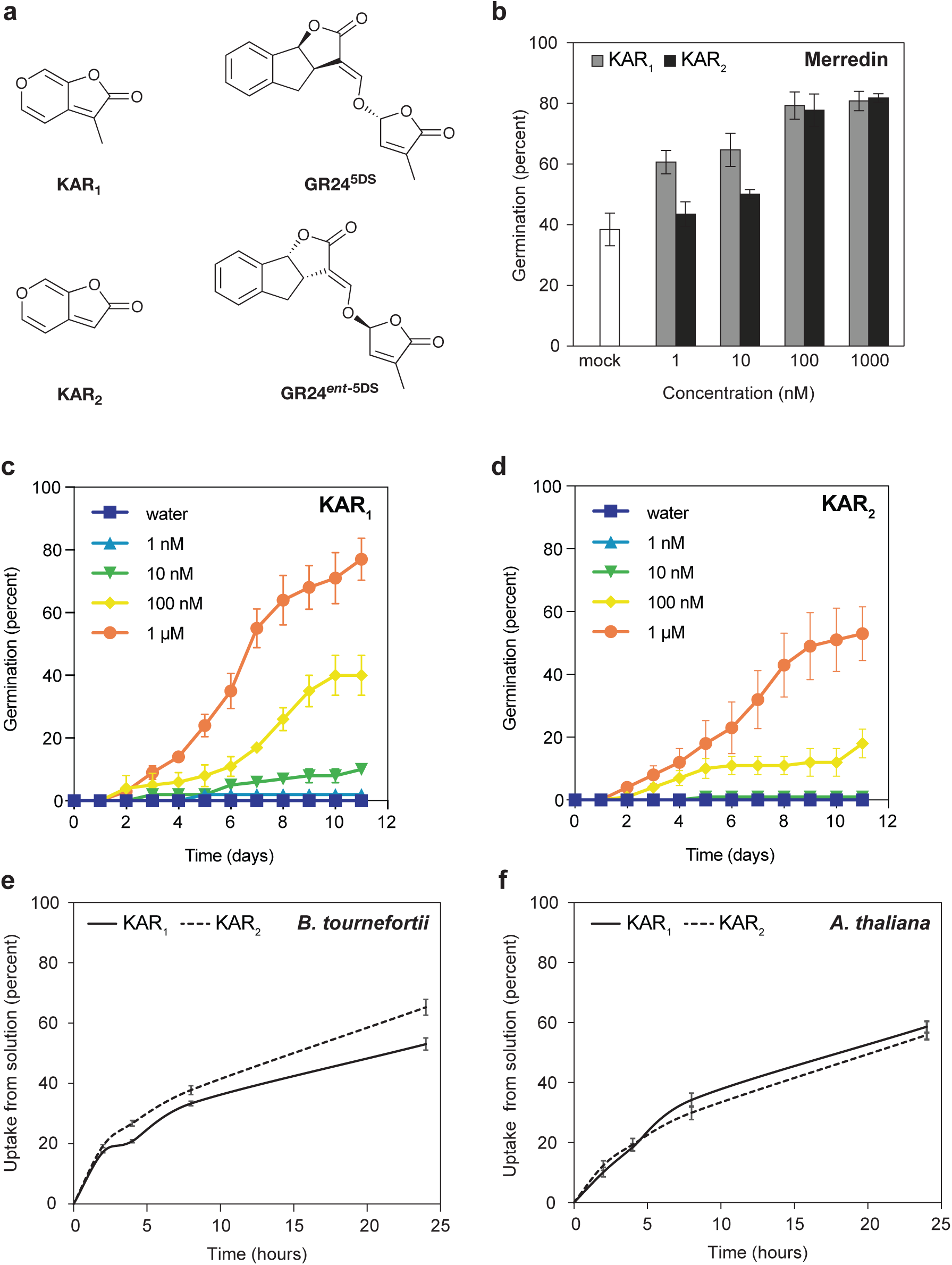
Germination of *Brassica tournefortii* seed treated with karrikins. **a**, Structures of KAR_1_, KAR_2_ and two enantiomers of GR24. b, Germination response of the “Merridin” batch of *B. tournefortii* seed treated with KAR_1_ and KAR_2_ after three days. c-d, Germination response of the “Perth” batch to KAR_1_ (c) and KAR_2_ (d). Data in Figure 1a are derived from the data shown in c. e-f, Uptake of KAR_1_ and KAR_2_ by imbibed seed, as determined by GC-MS. All error bars are mean ± SE of n = 3 batches of 75 seed (b-d) or 3 samples of 40 mg (e) or 20 mg (f) seed as described in Methods. Source data are provided as a Source Data file.

**Supplementary Figure 2.**
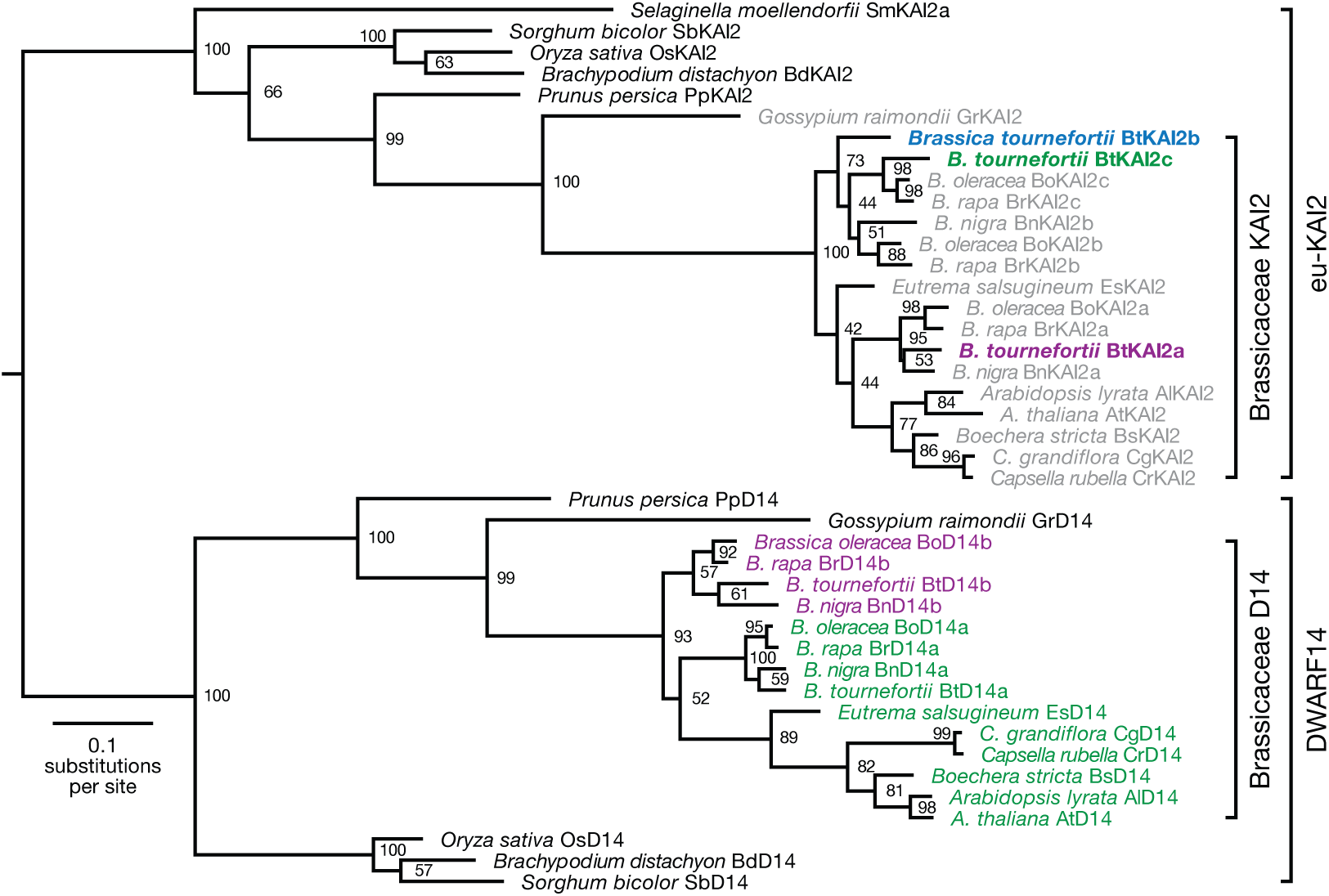
Extended phylogeny of eu-KAI2 and D14 proteins in angiosperms. Maximum likelihood phylogeny of KAI2 and D14 homologues in the Brassicaceae and monocots, based on nucleotide data. Node values represent bootstrap support from 100 replicates. A KAI2 sequence from *Selaginella moellendorffii* (SmKAI2a) serves as an outgroup for the eu-KAI2 clade. Source data are provided as a Source Data file.

**Supplementary Figure 3.**
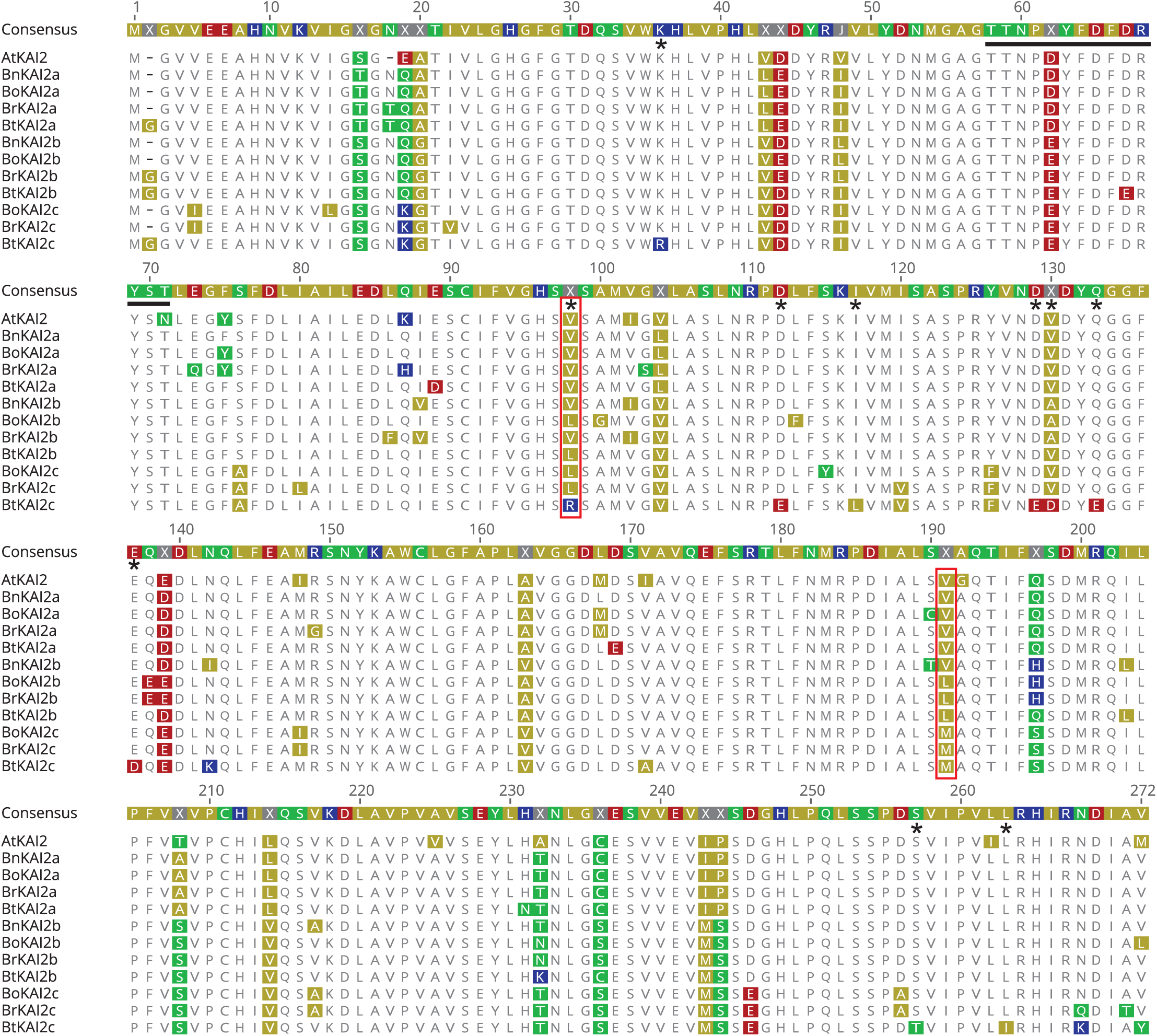
Alignment of Brassica KAI2 sequences. Full length protein coding regions of *KAI2* homologues from four *Brassica* species (*Brassica tournefortii*, *B. rapa*, *B. nigra* and *B. oleracea*) were translated from database nucleic acid sequences and aligned to *Arabidopsis thaliana* KAI2 using MAFFT^1^ implemented in Geneious R10 software (Biomatters Ltd). Amino acid residues are coloured according to polarity: yellow, non-polar (G, A, V, L, I, F, W, M, P); green, polar & uncharged (S, T, C, Y, N, Q); red, polar & acidic (D, E); blue, polar & basic (K, R, H). Residues that are unique to BtKAI2c but otherwise invariant are highlighted with asterisks (*); residues 98 and 191 are highlighted with red boxes. Source data are provided as a Source Data file.

**Supplementary Figure 4.**
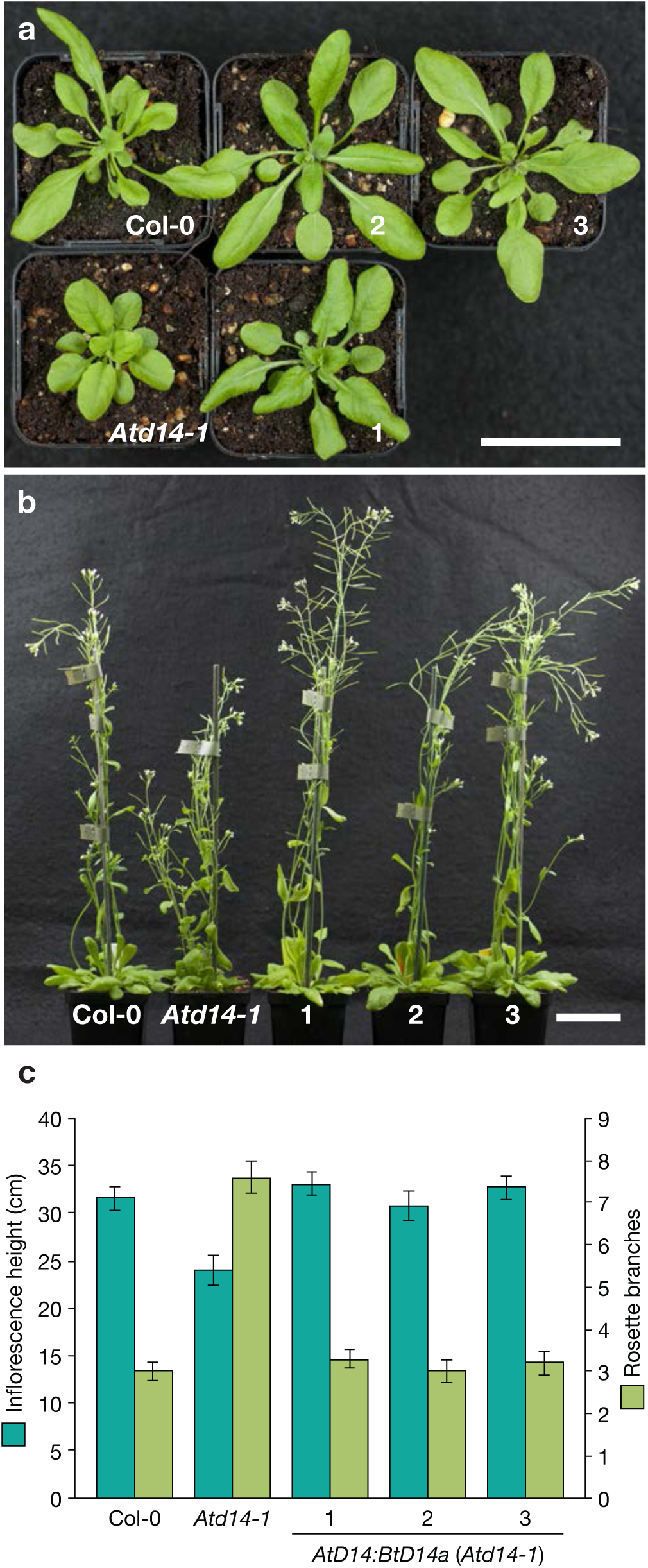
BtD14a is functionally homologous to AtD14. Three independent transgenic lines of Arabidopsis *Atd14-1* were analysed for functional complementation of the mutant phenotype by an *AtD14pro:BtD14a* transgene. a, Rosette and leaf morphology at 31 days post-germination. b, Plant height and number of primary rosette branches at 45 days post-germination. c, Quantification of height and branching parameters, *n* = 10 plants per genotype. Scale bars: 50 mm. Source data are provided as a Source Data file.

**Supplementary Figure 5.**
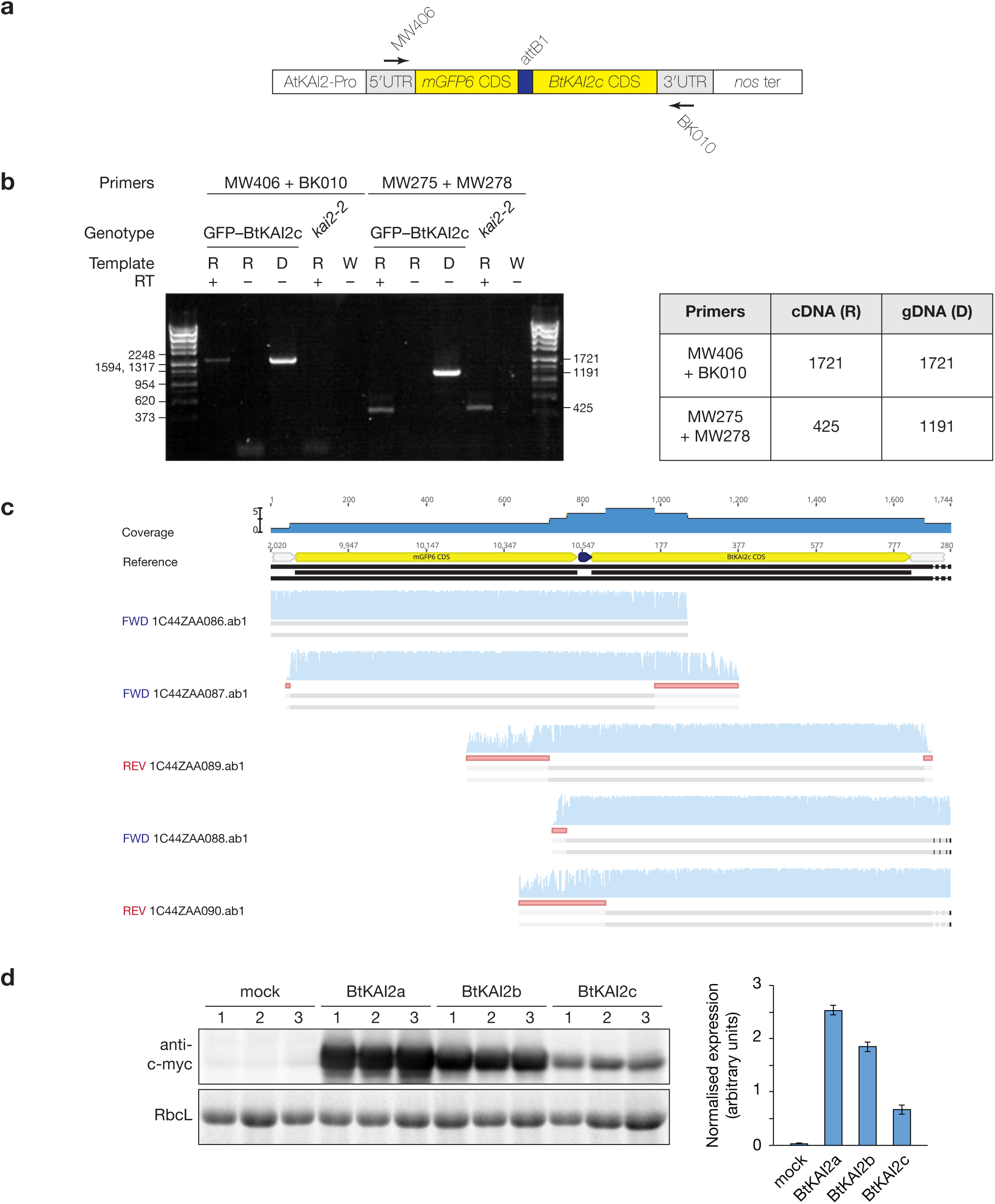
The GFP-BtKAI2c transgene is faithfully transcribed in Arabidopsis. **a**, Structure of the *AtKAI2pro:mGFP6-BtKAI2c* transgene. Primers used for RT-PCR are shown with arrows. The promoter and 5′UTR are derived from *At4g37470* (*AtKAI2*). attB1, Gateway recombination site that links *mGFP6* and *BtKAI2c* regions; *nos ter*, nopaline synthase terminator. Not drawn to scale. **b**, RT-PCR analysis of *GFP-BtKAI2c* transcripts after 35 cycles of amplification. Primers MW406 + BK010 target the transgene, while a second primer pair (MW275 + MW278) serves as a control and spans five introns of *At1g03055*. *kai2-2* serves as a non-transgenic control genotype. Templates: R, total RNA; D, genomic DNA; W, water only. RT, reverse transcriptase (Supercript III). DNA size standards (in base pairs) are indicated on the left, with anticipated PCR product sizes shown on the right and defined in the table. **c**, The RT-PCR product generated with proof-reading polymerase (Q5, New England Biolabs) and primers MW406 and BK010 was cloned into pCR4-TOPO (Life Technologies). Five dideoxy sequence traces were aligned against the *GFP-BtKAI2c* transgene reference. No disagreements with the reference sequence were observed. Red bars indicate trimmed regions of sequence traces to remove low quality data. **d**, Transient expression of BtKAI2a, BtKAI2b and BtKAI2c proteins in tobacco. Plasmids encoding N-terminal, c-myc-tagged proteins were transferred to Agrobacterium, and the resulting strains used to infiltrate tobacco leaves. After 96 h, samples were harvested in triplicate (two to three leaves per sample). Mock-treated leaves were transformed with a plasmid encoding a non-tagged protein. Sixty micrograms of total protein were separated by SDS-PAGE, blotted and challenged with anti-c-myc antibody (Genscript A00704). Band intensity was measured using ImageJ, and expression was normalised to intensity of the Rubisco large subunit (RbcL) band on the “stain-free” gel imaged under UV light. Error bars indicate SE, n = 3 replicates. Source data are provided as a Source Data file.

**Supplementary Figure 6.**
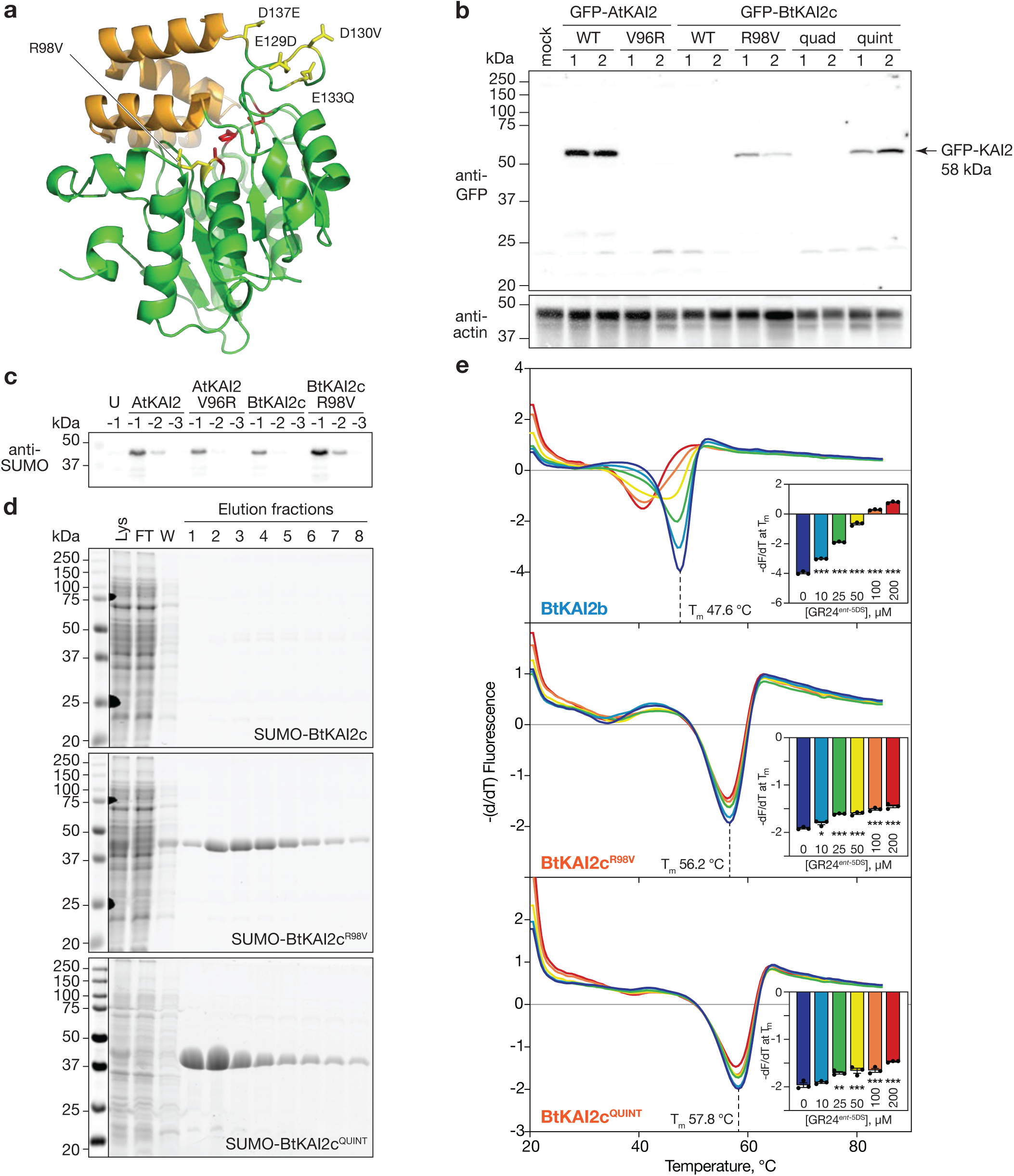
The R98V mutation restores stability of BtKAI2c. **a**, Homology model of the native BtKAI2c protein. Highlighted in yellow are five residues and their corresponding mutations that were assessed for effects on BtKAI2c stability. The combination of E129D, D130V, E133Q and D173E in the hinge region between the lid domain (orange) and core domain (green) is the quadruple (“quad”) mutant; combining this with R98V yields the quintuple (“quint”) mutant. Red residues are the Ser-His-Asp catalytic triad. b, Transient expression of AtKAI2 and BtKAI2c variants in tobacco leaves following infiltration by *Agrobacterium* strains encoding GFP-KAI2 variants driven by the CaMV *35S* promoter. Three leaves were infiltrated on each of two different plants. After five days, 20 µg soluble protein was electrophoresed, blotted and challenged with antibodies against either anti-GFP (top) or anti-actin (bottom). c, Immunoblot of crude bacterial lysates expressing SUMO fusion proteins and challenged with anti-SUMO antibody. Negative values above each lane indicate log10 dilution factors of lysate; U, lysate from uninduced bacterial culture expressing SUMO-AtKAI2. d, SDS-PAGE analysis of protein purification runs of SUMO-BtKAI2c, -BtKAI2c^R98V^ and -BtKAI2c^QUINT^ harvested from 900 mL bacterial culture. Lys, clarified lysate; FT, column flow through after immobilisation on Co^2+^ affinity column; W, wash with 10 mM imidazole. Proteins were eluted with 200 mM imidazole in eight successive fractions. e, DSF curves of SUMO fusion proteins treated with 0-200 µM GR24*^ent^*^-5DS^. Insets plot the minimum value of – (dF/dT) at the melting point of the protein as determined in the absence of ligand (means ± SE, n = 3). Significant differences from untreated control: * P < 0.05 ** P < 0.01 *** P < 0.001 (ANOVA). Source data are provided as a Source Data file.

**Supplementary Figure 7.**
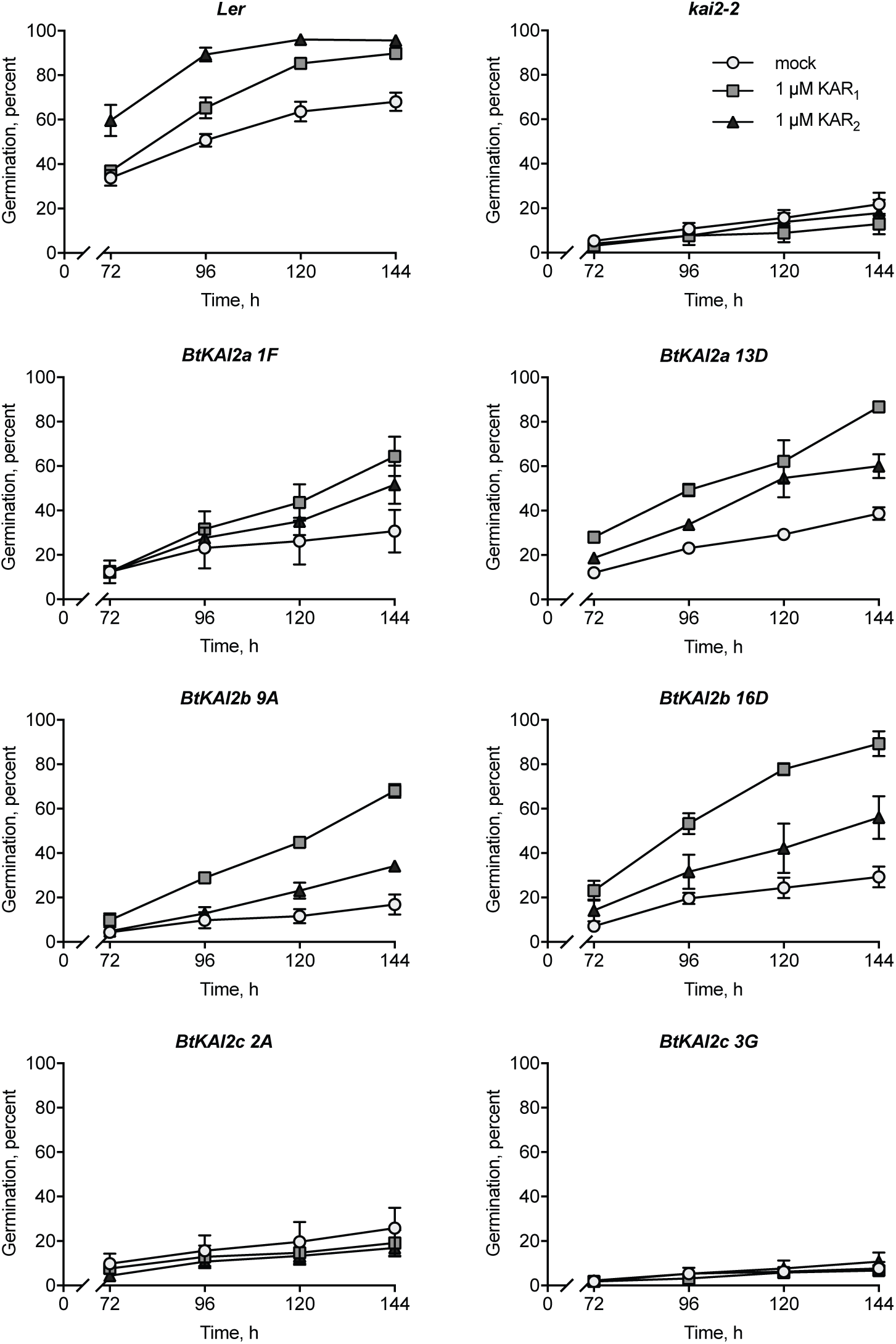
Germination profiles of transgenic Arabidopsis seeds expressing BtKAI2 homologues. Freshly harvested seed (three batches per genotype, each batch harvested from four plants) were removed from freezer storage, surface-sterilised and sown on 1% Phytagel supplemented with 0.1% acetone (mock), 1 µM KAR_1_ or 1 µM KAR_2_. Seed were incubated under constant light at 25 °C. Seed were examined for germination (radicle protrusion) 72 h after sowing and every 24 h thereafter. Data are means ± SE of three independent seed batches and 75 seed per batch. For each transgene, two independent, homozygous transgenic lines were analysed. Data presented in Figure 3 of the main manuscript are derived from these data. Source data are provided as a Source Data file.

**Supplementary Figure 8.**
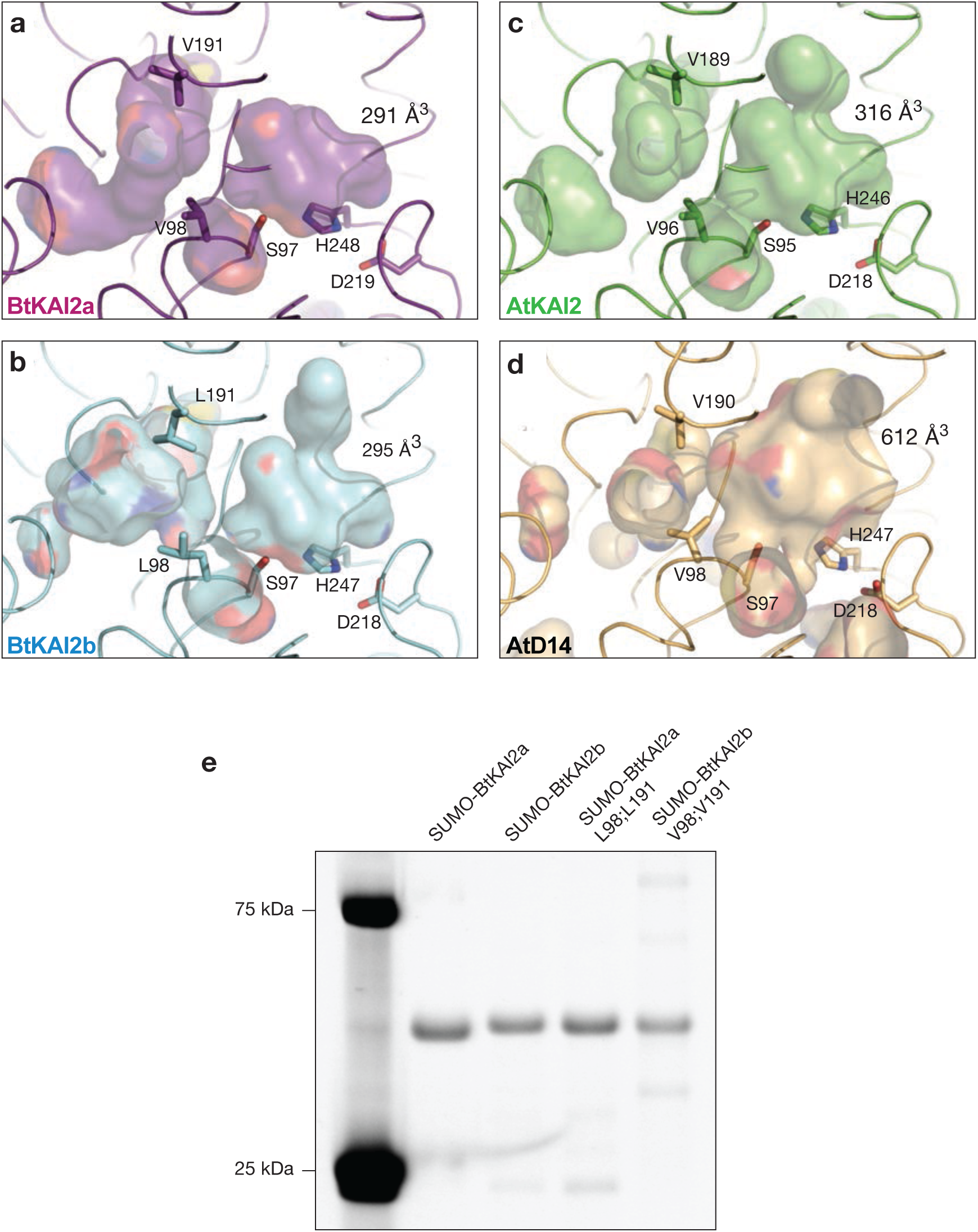
BtKAI2 homology models and SDS-PAGE of SUMO-BtKAI2 fusion proteins used for DSF. **a-d**, Solved structure of AtKAI2 (PDB: 3w06, ref. 2), AtD14 (PDB: 4IH4, ref. 3) and predicted homology models of BtKAI2a and BtKAI2b. Coloured surfaces depict internal cavities; values indicate the volumes of the primary ligand-binding cavities, adjacent to the catalytic Ser-His-Asp residues. Also shown is a variable secondary pocket, to the left of the primary pocket in these images. **e**, To assess purity after affinity chromatography, five micrograms of each purified protein was electrophoresed on a 12% acrylamide gel containing 2,2,2-trichloroethanol and visualised under UV light. Protein size standards at 75 and 25 kDa (Bio-Rad Precision Plus Dual Colour) fluoresce strongly under UV light. Source data are provided as a Source Data file.

**Supplementary Figure 9.**
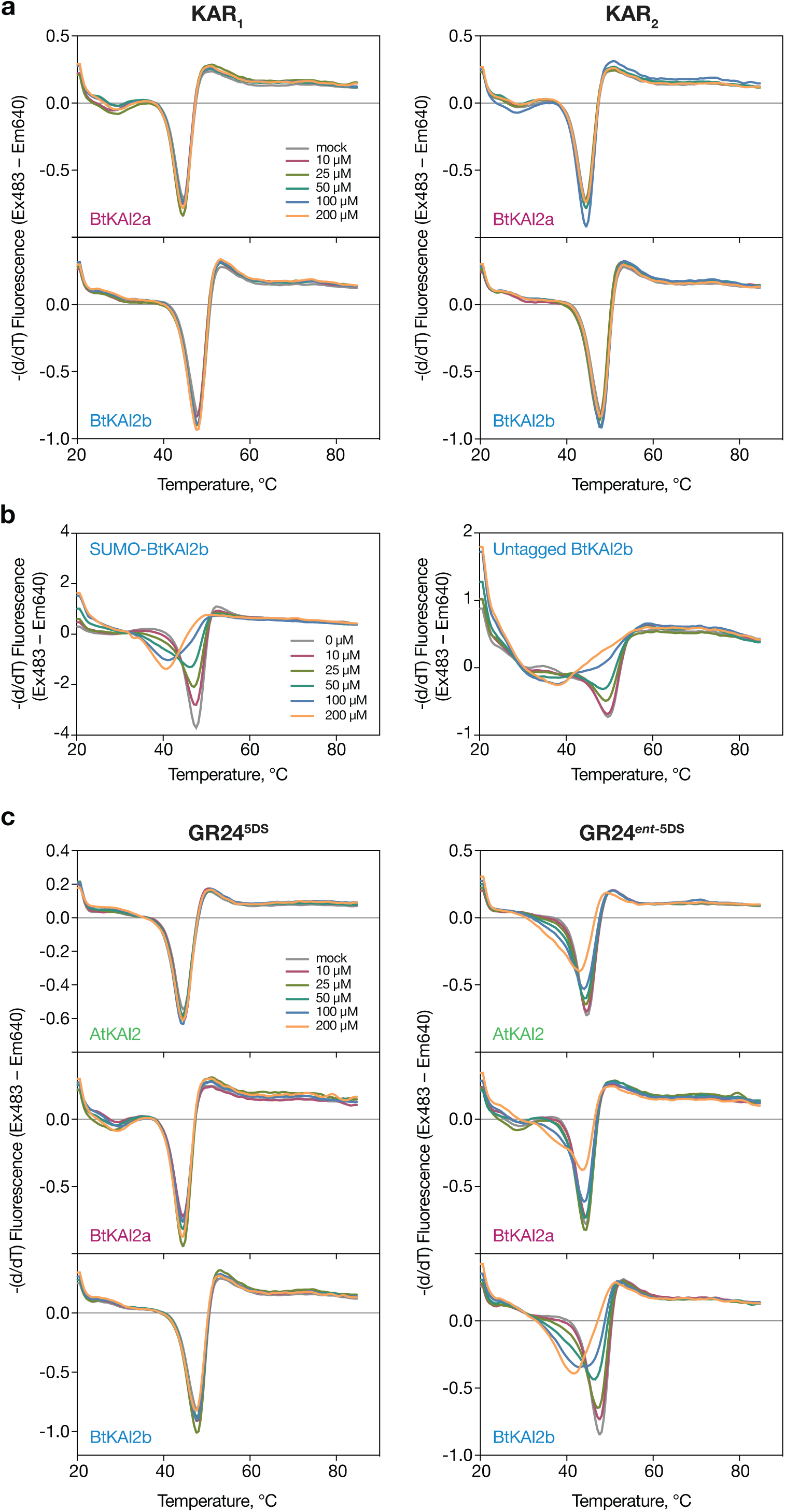
BtKAI2a and BtKAI2b do not respond to karrikins in DSF assays. **a**, Differential scanning fluorimetry curves of SUMO-BtKAI2a and SUMO-BtKAI2b in presence of 0–200 µM KAR_1_ or KAR_2_. b, DSF responses of SUMO-tagged BtKAI2b (left) or untagged BtKAI2b (right) to 0–200 µM GR24*^ent^*^-5DS^. c, DSF responses to 0–200 µM of the two enantiomers of GR24. Data are means of eight (a and c) or four (b) technical replicates at each concentration of ligand. Source data are provided as a Source Data file.

**Supplementary Figure 10.**
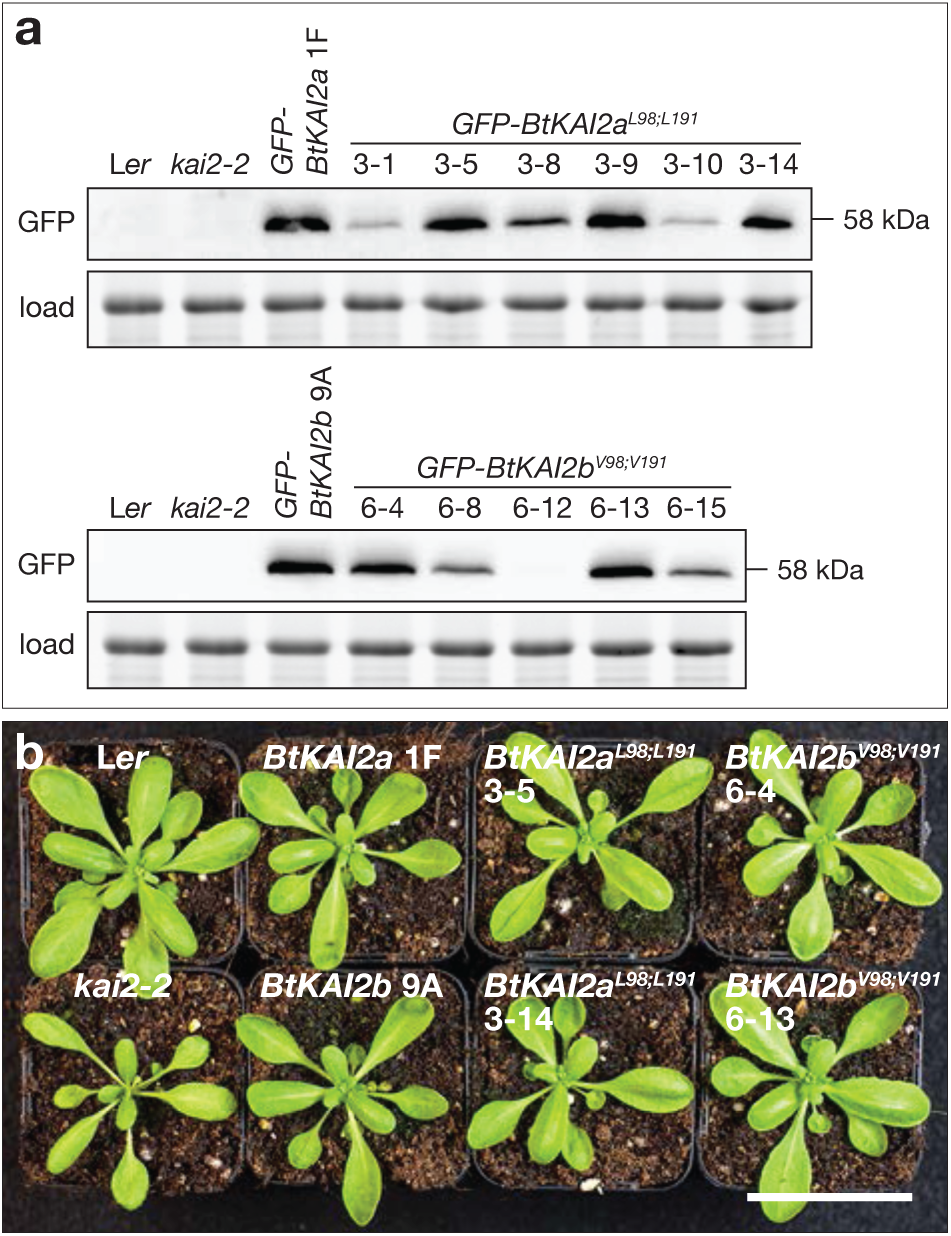
Stable transgenic expression of BtKAI2 valine–leucine exchange proteins in Arabidopsis. **a**, Immunoblots of total soluble protein extracted from 7-day-old seedlings of independent transgenic lines segregating in a 3:1 ratio for hygromycin resistance. Transgene expression in six lines expressing *GFP-BtKAI2a^L98;L191^* (upper panels) and five expressing *GFP-BtKAI2b^V98;V191^* (lower panels) were compared to a representative unmodified control (*GFP-BtKAI2a* 1F and *GFP-BtKAI2b* 9A respectively). Based on expression level two lines of each construct (3-5 and 3-14; 6-4 and 6-13) were selected and brought to homozygosity for further experiments. Protein blots were challenged with anti-GFP antibody. Equal gel loading was assessed by imaging total protein prior to blotting; the RbcL band is shown. b, Rosette phenotypes of homozygous individuals expressing native and modified *GFP-BtKAI2* transgenes. Plants were 25 days old and grown under long day conditions as described in Methods. Scale bar: 50 mm. Source data are provided as a Source Data file.

**Supplementary Figure 11.**
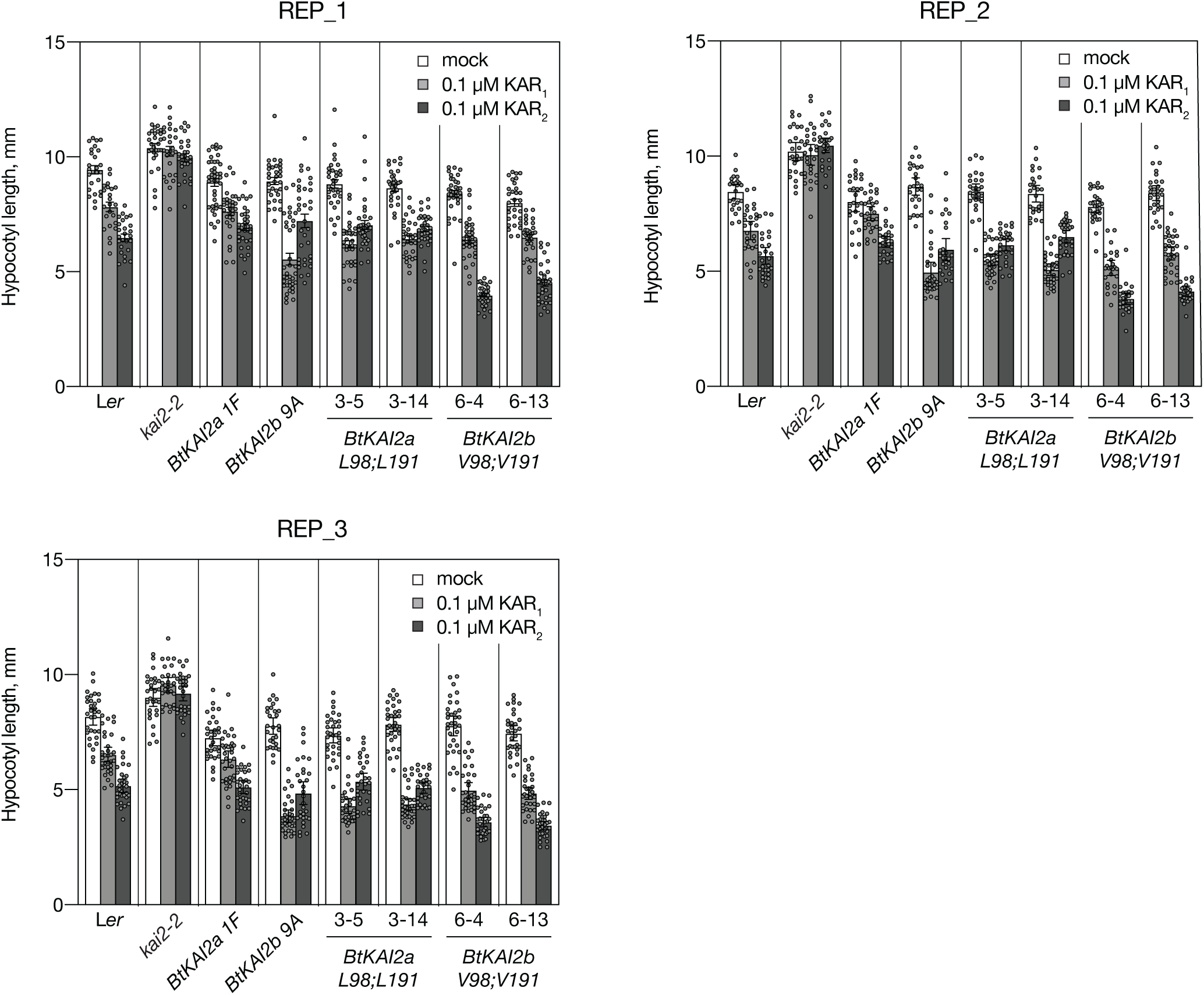
Three experimental replicates of hypocotyl elongation assays with BtKAI2a and BtKAI2b transgenics (double exchange of residues 98 & 191) Each panel depicts data from an independent experiment performed on a separate date, which are shown in summarised format in Figure 4. Data are means ± SE, n=24 to 40 seedlings. Each dot corresponds to an individual seedling. Source data are provided as a Source Data file.

**Supplementary Figure 12.**
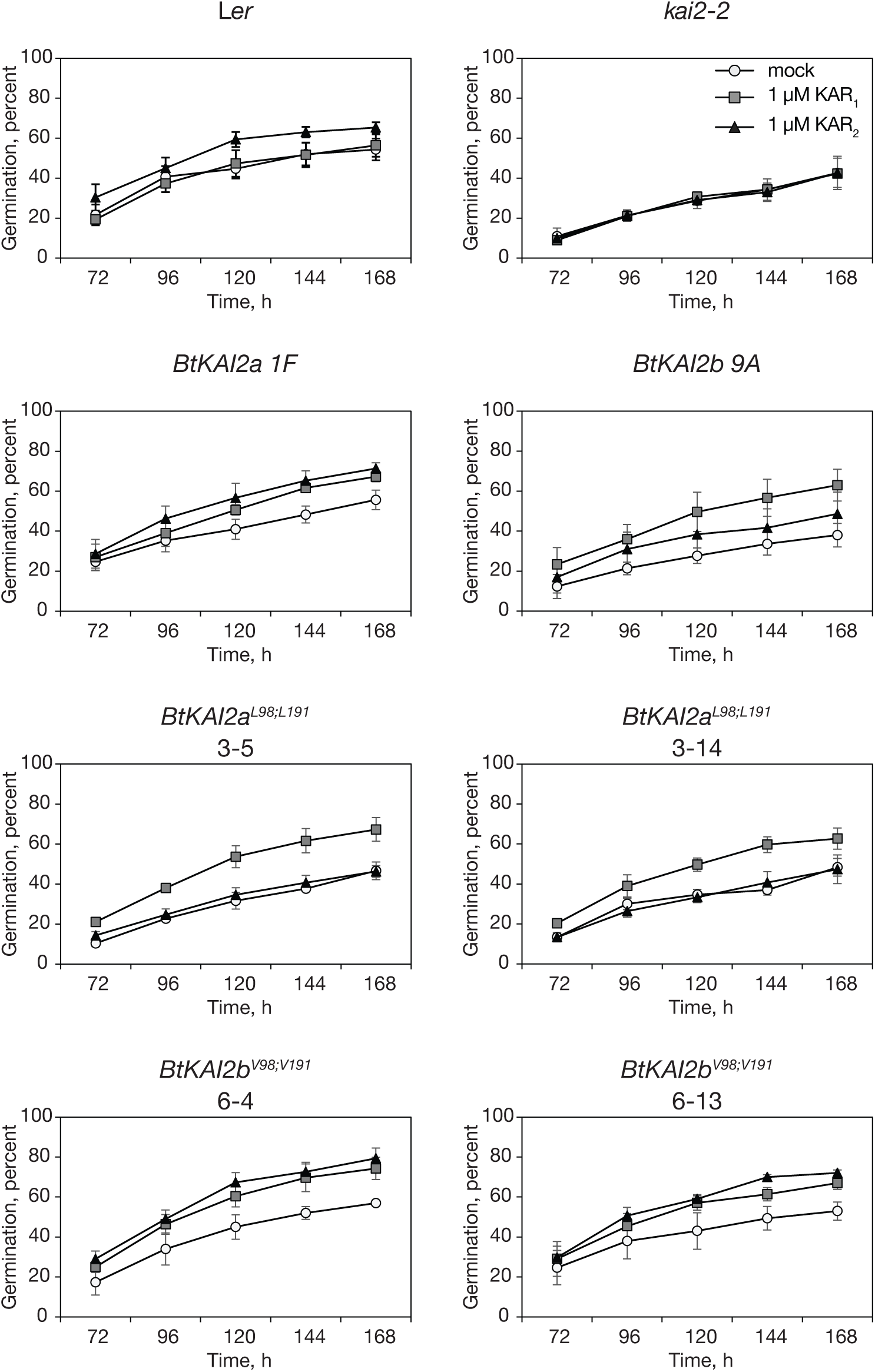
Germination profiles of transgenic Arabidopsis seeds expressing BtKAI2 homologues (double exchange of residues 98 and 191) Freshly harvested seeds (three batches per genotype, each batch harvested from three plants) were removed from freezer storage, surface-sterilised and sown on 0.7% agar supplemented with 0.02% acetone (mock), 1 µM KAR_1_ or 1 µM KAR_2_. Seeds were incubated under constant light at 25 °C. Seeds were examined for germination (radicle protrusion) 72 h after sowing and every 24 h thereafter. Data are means ± SE of three independent seed batches and 100 seed per batch. For the *BtKAI2a^L98;L191^* and *BtKAI2b^V98;V191^* transgenes, two independent, homozygous transgenic lines were analysed. Source data are provided as a Source Data file.

**Supplementary Figure 13.**
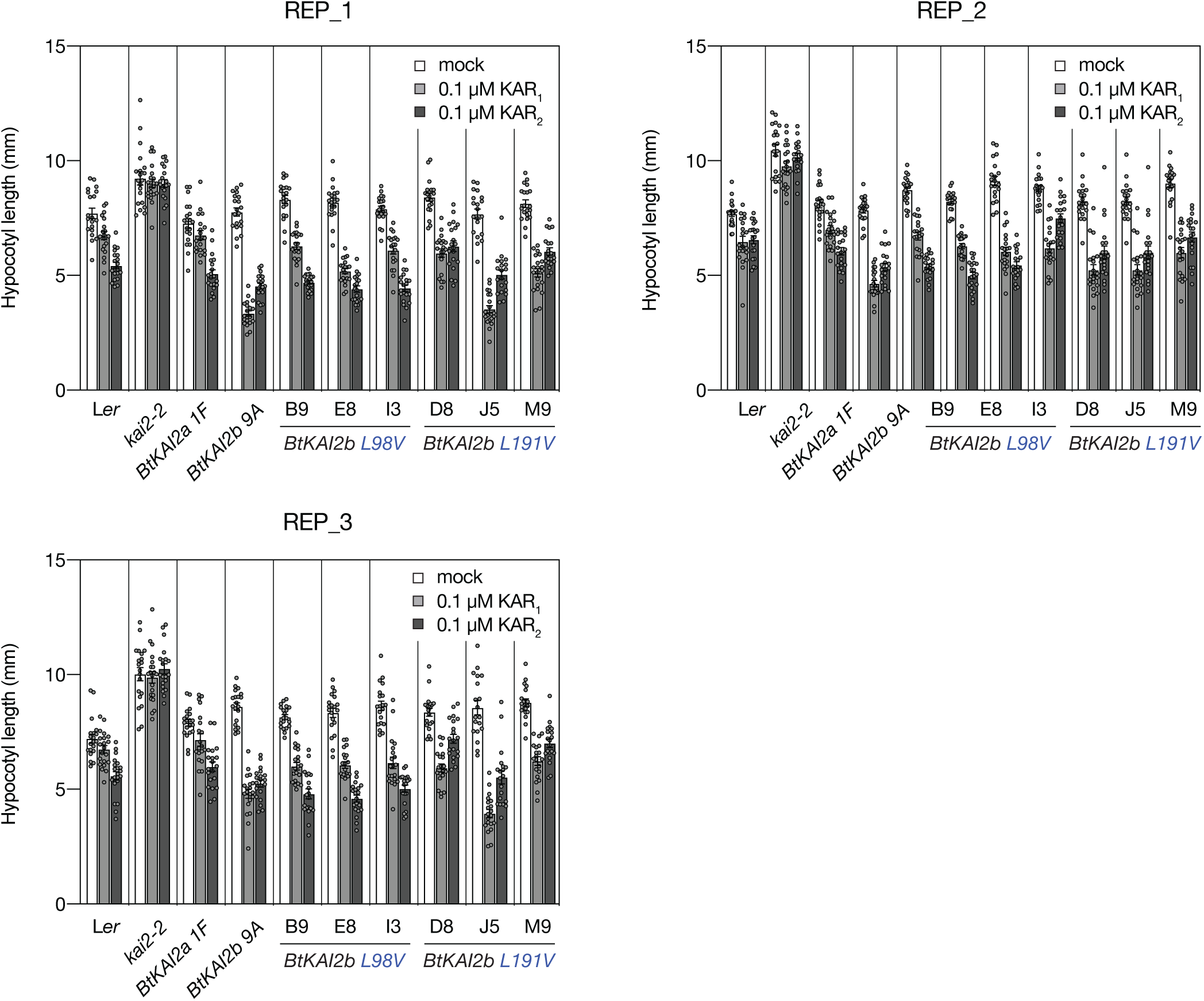
Three experimental replicates of hypocotyl elongation assays with BtKAI2b transgenics (individual exchange of residues 98 and 191) Each panel depicts data from a separate experiment performed on the indicated date, which are shown in summarised format in Figure 5. Data are means ± SE, n = 19-21 seedlings. Each dot corresponds to an individual seedling. Source data are provided as a Source Data file.

